# High coverage of single cell genomes by T7-assisted enzymatic methyl-sequencing

**DOI:** 10.1101/2022.02.23.481567

**Authors:** Juan Wang, Yitong Fang, WenFang Chen, Chen Zhang, Zhichao Chen, Zhe Xie, Zhe Weng, Weitian Chen, Fengying Ruan, Yeming Xie, Yuxin Sun, Mei Guo, Yaning Li, Chong Tang

## Abstract

Conventional approaches to studying 5mC marks in single cells or samples with picogram input DNA amounts usually suffer from low genome coverage due to DNA degradation. Many methods have been developed to optimize the library construction efficiency for bisulfite-treated DNA. However, most of these approaches ignored the amplification bias of bisulfite-treated DNA, which leads to shallow genome coverage. In this study, we developed the T7-assisted enzymatic methyl-sequencing method (TEAM-seq), which adopts enzymatic conversion to minimize DNA degradation and T7 polymerase-assisted unbiased amplification. We demonstrate that TEAM-seq delivered, to the best of our knowledge, the highest reported coverage(70% for 100pg, 35% for 20pg) of single cell genomes in whole-genome 5mC sequencing.

## Introduction

In recent years, the rapid development of single-cell sequencing technology has provided many valuable insights into complex biological systems, for example, revealing complex and rare cell populations and tracking the trajectories of distinct cell lineages in development ^1^. Numerous single-cell RNA and DNA sequencing methods have been developed ^2^. Heritable phenotype changes occur not only owing to changes in DNA and RNA nucleotide sequence, but also via epigenetic modifications, which do not change the DNA sequence itself. Among the epigenetic modifications, methylation of cytosine with the formation of 5-methylcytosine (5mC) is the most abundant epigenetic base change in vertebrates. This epigenetic modification has been intensively studied in relation to embryonic development, genomic imprinting, X-inactivation, cellular proliferation, and differentiation ^3, 4^. Conventional approaches to studying 5mC epigenetic marks usually rely on chemical treatment to convert modified/unmodified sites to readable mutated nucleotides, which requires high DNA input ^5^. However, the limited number of DNA copies significantly complicates the application of these conventional approaches to studies in single cells.

The standard and well-established technology to detect sequence-specific 5mC marks is bisulfite sequencing ^6, 7^, which utilizes sodium bisulfite to convert the unmethylated cytosine to uracil after constructing the sequencing library. Methylated cytosine is resistant to bisulfite conversion. Although bisulfite sequencing has been established as the gold standard for bulk DNA methylation analysis, single-cell adaptations of this method for low DNA input face a major hurdle of bisulfite-induced DNA degradation. Pioneer studies of single-cell bisulfite sequencing were performed by integrating all conventional steps, including bisulfite sequencing, into a single-tube reaction, followed by PCR amplification and deep sequencing ^6, 7^. The constructed DNA library is usually broken by the harsh bisulfite treatment, and the CpG/genome coverage drops to 5%/4% ^8^. The CpG coverage rate can be improved to approximately 18% through post-bisulfite adaptor tagging ^9^. Single-cell genome-wide bisulfite sequencing, in which the adaptor tags bisulfite-treated DNA with a 3′ stretch of random nucleotides was shown to further improve the coverage by up to 5–48.4% of all CpG sites ^10–13^. Recent advances in single-nucleus methylcytosine sequencing have been based on this approach, with an additional use of adaptase and random priming to tag bisulfite-treated DNA with adaptors ^14^.

In addition, recent advances have facilitated high-throughput single-cell methylation sequencing. Single-cell combinatorial indexing for methylation analysis uses transposons to assign combinatorial indices on single cells and then tag the other adaptor by random priming after bisulfite conversion ^15^. Although these previous methods improved the adaptor tailing efficiency, they suffered from DNA degradation because of the harsh bisulfite treatment. Recently, a gentle conversion method that combines ten-eleven translocation (TET) oxidation with pyridine borane sequencing was reported ^16^, which used a high concentration of TET1 to convert 5mC to 5CaC, which in turn, was converted to uracil by bicarbonate treatment. However, high concentrations of TET1 are not available in most laboratories. Overall, single-cell methylation sequencing coverage was poor, as it covered only 20% of CpG sites and less than 10% of the genome. Therefore, it is necessary to improve the coverage, accuracy, and read length of single-cell methylation sequencing.

The methods mentioned above improve the adaptor tagging efficiency or minimize DNA loss, but they ignore biased PCR amplification that causes uniform genome coverage. Here, we describe the T7-assisted enzymatic methyl-sequencing method (TEAM-seq) that improves coverage. In TEAM-seq, transposons tagmentate genomic DNA with T7 adaptors. Then, a gentle and high conversion efficiency kit is used to protect the methylation sites by TET2 and convert the unmethylated sites to uracil by the activity of apolipoprotein B mRNA editing enzyme catalytic subunit 3A (APOEC3A). This method can also be easily scaled up to high-throughput analysis of single cells. With TEAM-seq, we achieved a superior 70% coverage efficiency with only 2× sequencing depth and ultra-low DNA input, which, to the best of our knowledge, is the highest single-cell methylation coverage ever reported.

## Results

### Feasibility of enzyme-based conversion to detect the methylome

To detect the methylome, the conventional method is bisulfite sequencing (BS), which relies on the bisulfite-induced conversion of cytidine to uridine in the genome. However, 5-methylcytidine (5mC) does not get converted in this way (Sfig.1a). Furthermore, this harsh treatment usually results in very short fragments and significant loss of genomic DNA ^17^. To avoid the harsh bisulfite treatment, we consider using the gentle enzyme-based conversion methods, such as TET-assisted pyridine borane sequencing (TAPS)^16^ or commercially available EM-seq (NEBNext® Enzymatic Methyl-seq Kit | NEB). Unlike directly converting 5mC to U by TAPS, the commercially available enzyme-based method termed Enzymatic Methyl-seq (EM-seq) has been developed recently by New England Biolabs, which uses APOEC3A to convert cytidine to uridine for a gentler treatment of DNA ^18^. The conversion protocol included two general steps: 1) TET2 transformed 5mC/5hmC to 5-formylcytosine (5fC), 5-carboxycytosine (5caC), and 5-(β-glucosyloxymethyl)cytosine (5gmC), which could be protected from downstream APOEC3A conversion; 2) APOEC3A converted all the remaining unmodified cytidines to uridines (Sfig.1a). The unconverted “C” in sequencing data represented methylation sites, as in BS. A comprehensive analysis of the catalytic activity of this enzyme and a comparative study of the NA12878 dataset have been published previously^19^. Here, we sought to further extend the application of EM-seq to plant genomic DNA of *Arabidopsis*.

The BS and EM-seq libraries were prepared in a pairwise manner. We used 1μg of *Arabidopsis* DNA for the initial test. Both BS and EM-seq require PhiXDNA to balance the sequencing signal bias. The sequencing quality and data yield were comparable between EM-seq and BS (Sfig1.b–f). EM-seq was characterized by a **14%** lower duplication rate (Sfig.1c) and better methylome coverage (Sfig.1d, e) compared to those observed with the BS approach. To assess the accuracy of EM-seq, we added two different types of DNA to estimate the false-positive rate (lambda DNA) and 600 bp synthetic methylated DNA to estimate the false-negative rate. We found that false positive and false negative rates were comparable for EM-seq and BS approaches (1.69% vs. 0.96% and 6.03% vs. 2.93%, respectively) (Sfig.1c). The conversion rate of unmodified C in EM-seq was approximately **98.31%** (Sfig.1c). The same software and pipeline were used to analyze the outputs of BS and EM-seq experiments (Supplementalmaterial1_detailedprotocol).

Next, we analyzed the consistency of EM-seq and BS methylome data. At least four reads covered more than 15 million CpG positions in both methods. We defined a base as methylated if its methylation ratio was greater than 0.5 and found that 98.5% of CpG regions (covered > 4 times) had a consistent modification pattern in EM-seq and BS experiments. (Fig. 1a). For example, both EM-seq and BS reported highly correlated methylation sites across chromosomal regions (Fig. 1b). By further comparing the modification level for each 5mC covered by at least **three** reads, we observed a good correlation (0.89) between EM-seq and BS data (Fig. 1c). The density of the methylation across chromosome 4 or on gene elements were similar between the two methods (Fig. 1d–f). For example, both methods showed equal methylation level distribution around the CpG islands (CGIs) (Fig. 1g). Together, these results indicate that EM-seq can directly replace whole genome BS (WGBS) and provide comparable results. We then further examined whether this non-destructive method could be used for methylation detection in ultra-low input samples, using various conversion methods.

**Fig 1.**
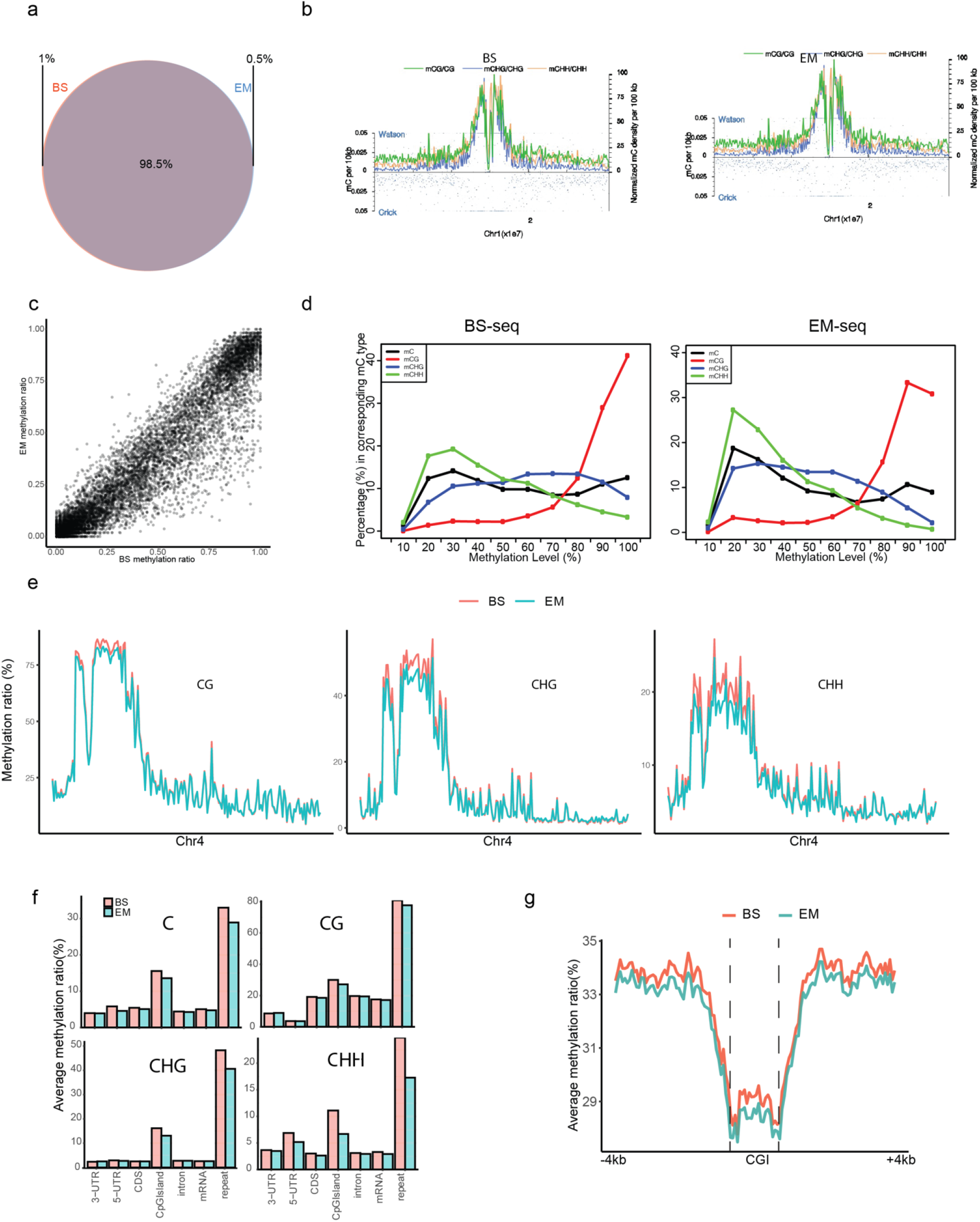
The data consistency between BS-seq and EM-seq. We annotated the confident methylation sites with at least four reads covered and the methylation ratio is larger than 0.5. 98.5% methylated sites in BS-seq were detected in both EM-seq and BS-seq (a). We used the Chr1 as an example, and we found that the methylated CG/CHH/CHG sites identified by these two methods were distributed similarly across the Chr1 (b). We further looked at the methylation level on each site. The global methylation ratio correlation is around 0.89 (spearman, *p*<0.05) (c). The methylation ratio density plot also showed the similar pattern between two methods. The medium methylation level of CHH/CHG in EM-seq was slightly lower than in the BS-seq (d). For example, the CG/CHG/CHH methylation ratio distribution across Chr4 were similar between EM-seq and BS-seq. The CHH/CHG methylation level in EM-seq demonstrated slightly lower level than in BS-seq in highly methylated regions (5% lower) (e). By further looking at the gene elements, the average methylation ratios on the gene elements were also slightly lower in EM-seq than in BS-seq, typically for CHH (f). More detailed investigations on methylation patterns around CpG islands (CGIs) showed the methylation pattern distributed similarly by using these two methods (g).

### DNA polymerase amplification of a large uridine-rich fragment

Multiple displacement amplification (MDA) is one of the most popular choices for non-destructive amplification of the genome in ultra-low input samples ^20, 21^. We started by performing EM-MDA-seq in a 1-ng sample of genomic DNA (Fig. 2b). Genome cytidines were converted to uridines by EM-seq, producing long uridine-rich genomic fragments (Fig. 2a). The converted DNA fragments underwent MDA and 10-fold amplification was achieved after 30 min of incubation (Fig. 2b). Subsequently, conventional DNA library construction was performed, and EM-MDA-seq of this 1-ng input showed excellent performance (**84.79%** coverage at the **6.96×** depth) (Fig. 2c, green). However, in the experiment with a picogram input, EM-MDA-seq showed a large amplification bias and insufficient coverage (**5.41%** coverage at the **2.62×** depth) (Fig. 2c, green). We then decided to lower the EM-MDA-seq bias with picogram input samples by improving the priming efficiency and polymerase speed.

**Fig 2.**
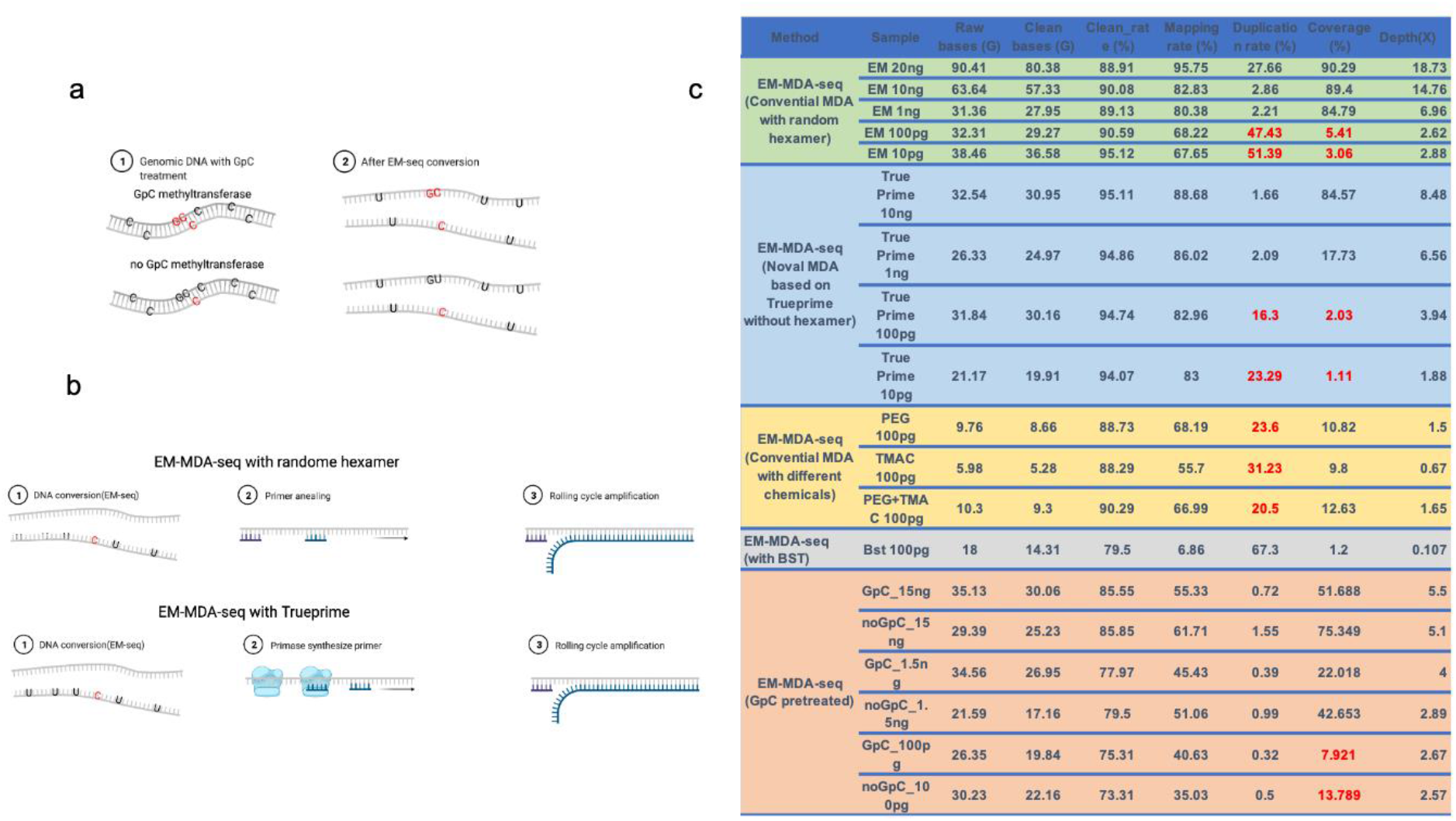
The MDA amplification on the EM-seq converted genome. After EM-seq conversion, most of the unmodified Cs were converted to U by APOEC3A deaminase. The GC content was decreased nearly 48%. The methylated Cs were protected by Tet2/enhancer, and stay as the cytidine analogs (5caC,5gmC,5fC) in the deamination process. The non-destructive method produced the pretty long uridine-rich ssDNAs (∼2kb). We also used GpC methyltransferase to pretreat the genomic DNAs. Considering the GpCs were rare on mammalian genome, the artificial GpCs should not affect the native methylation identification (CpG,CHH,CHG). The methylated GpC could also be preserved in the deamination step (a). The MDA method used the random hexamers to prime on the uridine-rich ssDNAs from EM-seq. Phi29 extended on the hexamers and displaced the DNA ahead. Then the displaced ssDNA could be amplified again by random hexamers. The process was termed as “EM-MDA-seq” (b). The hexamers annealing efficiency is largely rely on the GC contents and annealing temperature. To avoid the random hexamer annealing bias, the novel method Trueprime, the combination the TthPrimPol’s unique ability to synthesize DNA primers with the highly processive Phi29 DNA polymerase (Φ29DNApol) enables near-complete whole genome amplification (b). The EM-MDA-seq of high input DNA (20∼1ng) produce the normal genome coverage around 80%. However, the EM-MDA-seq of picogram input DNA only give us 3∼5% genome coverage with very high duplication rate 47%. The Trueprime amplification also resulted the similar low genome coverage 1∼2% for picogram input DNA. The PEG have the weakly positive effect to improve the genome coverage, and TMAC did not show this sign either. We then tried use robust BST to replace Phi29 in MDA, which also produce the shallow genome coverage 1.2%. We then tried to use GpC methyltransferase to preserve more cytidine in the deamination step by converting all the GpCs to artificially methylated GpCs(a). By side-to-side comparison with the non-GpC treatment control, the GpC methyltransferase treatment did not demonstrated the significant advantages in genome coverage from nanogram input to picogram input. The methods were described in the supplementalmaterial1_detailedprotocol part D.

It has been previously reported that random hexamer priming results in a bias in nucleotide composition and that this bias influences the uniformity of the location of the reads. The novel MDA “Trueprime” method, which is based on the use of DNA primase without a random hexamer, demonstrated superior breadth and uniformity of genome coverage with high reproducibility ^22^ (Fig. 2b). However, Trueprime was not very efficient in amplifying the uridine-rich converted DNA and yielded only **2.03%** coverage at the **3.94×** depth. (Fig. 2c, blue).

It has been noted previously that DNA polymerase has base-pairing preference for nucleotide incorporation ^23^, which may cause amplification bias during EM-MDA-seq. Most DNA polymerases incorporate GT and AC nucleotides with high and low efficiency, respectively ^23^. In our experiments, we also found that the speed of Phi29 was 5-fold lower in bisulfite-converted DNA than in untreated genomic DNA (based on BGI production center reports), supporting the notion of the low efficiency of adenosine incorporation. We tried several solutions to normalize the nucleotide incorporation speed, including poly (ethylene glycol)/trimethylammonium chloride solution, slow down amplification, and minimizing the bias. We also tried an alternative robust polymerase, BST. However, none of these approaches showed superiority in terms of uniform coverage (Fig. 2c). However, to solve the issue of the AU bias in EM-seq, we used the GpC methyltransferase M.CviPI to pretreat the genome, which preserves 25% more GC content through the EM-seq conversion (Fig. 2a). Surprisingly, GpC methyltransferase treatment did not lead to obvious advantages over the control group in a side-to-side comparison (Fig. 2c orange).

Thus, the DNA polymerases Phi29 and BST had amplification bias in the converted uridine-rich DNA and were concluded to be unsuitable for EM-seq amplification. We then pursued an alternative amplification method with a more even coverage, by using T7 RNA polymerase amplification.

### Uniform amplification of ultra-low input uridine-rich DNA fragments by T7 RNA polymerase

Considering that the AC/GT bias amplification may be general problem for DNA polymerase, we used RNA polymerase instead of DNA polymerase to avoid the bias amplification. An improved single-cell whole genome amplification method based on the use of T7 polymerase has been recently reported that outperformed other MDA approaches, enabling a more uniform coverage of the genome ^24^. Unlike the MDA, using Phi29 DNA polymerase to ceaselessly synthesize the complementary strand on the single strand DNAs, T7 polymerase generated marvelous RNAs on the DNA templates with less bias. Inspired by that finding, we have developed a novel method, which we termed T7-Assisted Enzymatic methyl-seq (TEAM-seq).

In this method, we used Tn5 transposons to tagmentate the genome with biotinide adaptors and obtain medium-sized fragments (500–2,000 bp). The tagmentated DNA was then enzymatically converted, and single stranded DNAs were extended on the T7 promoter oligo to generate the T7 promoter on the DNA (Fig. 3a). Throughout the process, we used streptavidin beads to bind DNA without elution to minimize DNA loss in the purification. Finally, the enzymatically converted DNA was amplified using T7 RNA polymerase. The final constructed library was sequenced by the nanopore or Pacbio method to generate long reads for paternal/maternal phasing (the detailed protocol and experimental information can be found in the supplementarymaterial1). We used 100-pg or 20-pg of genomic DNA for the initial analysis. The coverage and other parameters were better or comparable to those achieved by the WGBS of a **1-μg** sample (**11.58%** duplication rate, coverage at **2.58×** depth). The consistency between TEAM-seq and WGBS approaches was approximately ∼93% (Fig. 3b). We also summarized the features of the different published methods (Fig.3c). We then performed a detailed study of the coverage and accuracy of TEAM-seq (comparing with scWGBS, and conventional WGBS).

**Fig 3.**
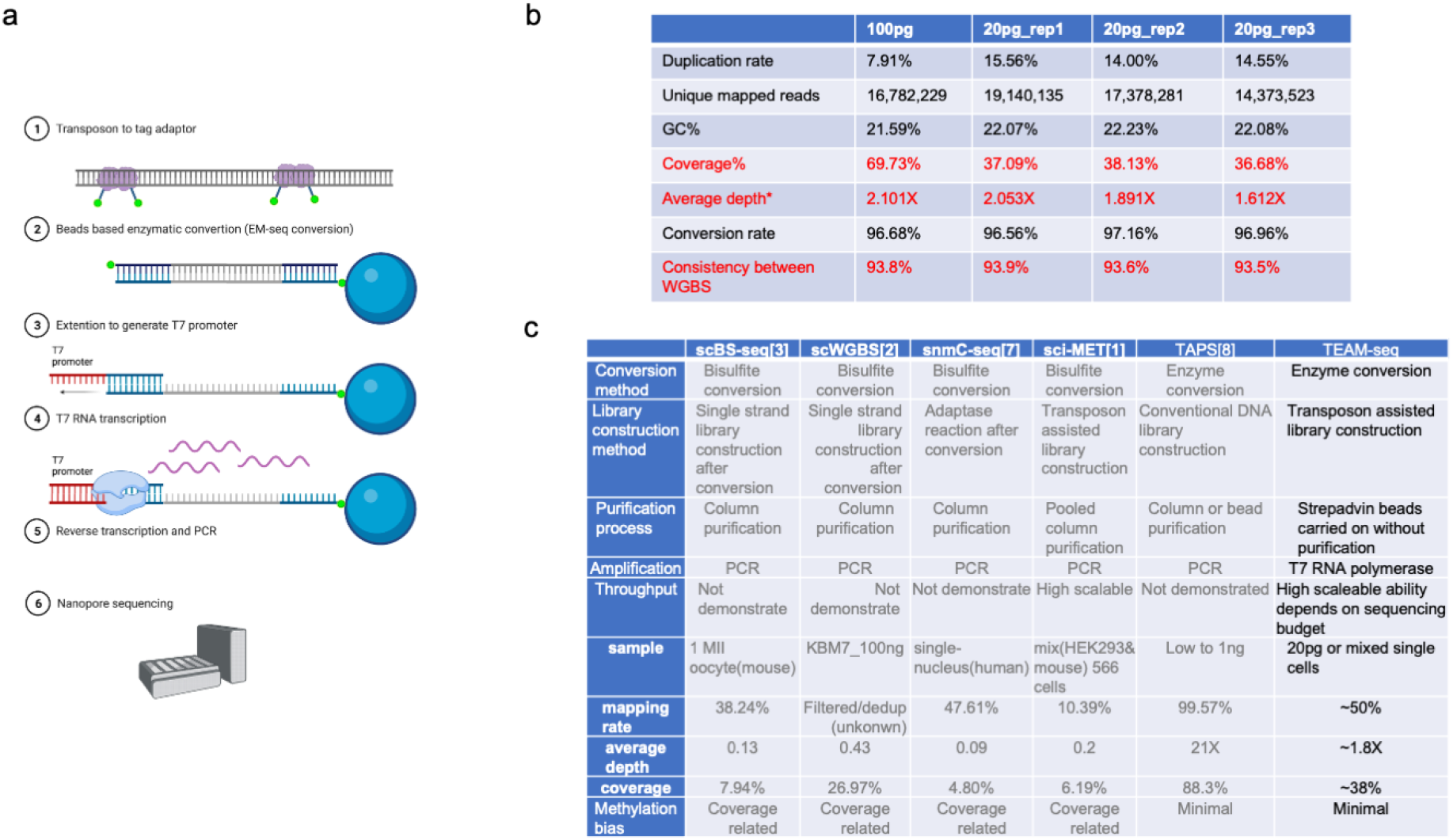
The schematic procedure of the TEAM-seq and its overall performance. The transposons were assembled with Tn5 enzyme and the DNA adaptor, with biotin on the on the 5’ end. After the tagmentation, the biotin adaptors were attached to the genomic DNAs and fragment it. After the denature the fragmented DNA were bound to the M280 strepadvin beads. Then the DNA carried on the beads went through the whole EM-seq protocol (detail could be found in method). Then the single stranded DNA were extended on the T7 oligos to generate the T7 promoters. After that, we performed the beads-on T7 transcription reaction for overnight. After the enough RNAs generated, the cDNA nanopore library were constructed and the detailed sequence information could be found on supplemental material_1. The final product could be sequenced on the nanopore(a). After we get the sequence, we analyze the data through the same protocol of the WGBS(1ug). The depth was calculated by average read count covered among all bases (ATGC). The whole genome coverage (the percentage of genomic sequence was covered by reads) is 69% for 100-pg TEAM-seq with 2x depth and 37% for 20-pg TEAM-seq with 1.8x depth. We selected high confidence sites to calculate the consistency. The methylated sites were defined as >0.5 methylation ratio with three covered reads, and the unmethylated sites were defined as <0.5 methylation ratio. The ∼93% genomic sites have the matched methylation status in the WGBS, showing the good consistency between low input TEAM-seq and the gold standard WGBS(b). (c) We summarized the genome coverages and key features of the different methods from previous publications ^1–4 5 6^. Column purification is the key step to limit the scalability. scBS-seq, scWGBS and snmC-seq constructed the library with barcodes after column purification. Therefore, the throughput of these three methods may be limited. The sci-MET and TEAM-seq constructed the library with barcodes before conversion, offering the simplicity in the downstream processing. The TEAM-seq took the advantages of the transposon library construction, beads carried on reaction and T7 RNA polymerase amplification to minimize the loss of the DNA and achieve uniform amplification.

The sequencing quality and data yield for 100-pg/20-pg samples were similar to those achieved by the conventional nanopore DNA sequencing. Analysis of the 20-pg input showed a higher base volatility than that of the larger, 100-pg input (Sfig. 2a). The 20-pg library resulted in **37.09%** genome coverage with **2.05×** sequencing depth, which are, to the best of our knowledge, the highest recorded values for such analyses (Fig. 3b). The accumulation curve showed the coverage saturation in each method (Sfig. 2b). For the picogram-level input, TEAM-seq showed lower GC bias and equal coverage of CGIs(CpG islands) than those achieved by the analyses of a 1-μg input by WGBS and 100-ng input by single cell WGBS (Sfig. 2c–e). The higher and more uniform genome coverage resulted in a larger number of all C sites covered by TEAM-seq in the case of a picogram input. In the experiment with a 20-pg input, TEAM-seq covered 26.97% of CpG and 24.75% of C sites at the **2×** depth, whereas for a 100 pg input, these values were higher: 54.85% and 52.79%, respectively. Further, TEAM-seq covered 34.12% and 63.74% of CGIs in experiments with 20 and 100-pg inputs, respectively.

TEAM-seq also showed a high consistency and methylation level correlation with the results of the gold standard WGBS. Over ∼**93%** of CpGs in TEAM-seq (100-pg/20-pg) showed modification states that matched those detected by WGBS (Sfig. 3a). The correlations between WGBS and TEAM-seq results (**100-pg**/**20-pg**) were **0.85/0.81** for the sites with >**20×** coverage. Distributions of the methylated sites were also very similar (average standard error ± 0.2 on CpG; ± 0.6 on CHG) on chromosome and gene element levels and close to those observed for WGBS (Sfig. 3c, d). The methylation ratios of the 20 pg genomic DNA sample or single-cell genomic DNA, which only had 1 or 2 genome copies, showed that the methylation distribution was polarized (peaks on 0 or 1) (Sfig. 3b). For example, in the 20 pg input sample, a similar methylation level distribution in the CGI was observed (Sfig. 3e). Moreover, we found that TEAM-seq had a shallow coverage-related methylation ratio bias (Sfig. 3f). Three technicians performed the experiments, and the reproducibility was sufficiently good (Sfig.4).

In summary, TEAM-seq uniformly amplified enzymatically converted DNA, resulting in better coverage of samples with ultra-low input DNA amount. Next, we used TEAM-seq to perform single-cell methylome analysis to expand its application scope.

### Indexed TEAM-seq for mid to high throughput single-cell analysis

Using transposons, we designed a unique, relatively long barcode, which could be identified by nanopore sequencing (Sfig. 5). After cell nuclei were sorted into plates, we labeled the cell genome using the barcoded transposons. We then performed single-cell TEAM-seq on the pooled cells (Please see Supplemental material 1_detailed protocol). Due to the throughput limitation of nanopore sequencing, we only processed 50–100 cells. Our proposed method can be easily scaled up and down based on your nanopore sequencing budget. We observed a meager collision rate of **5%**, based on the artificial mixture of the human(HeLa) and mouse(4T1) cells (Fig. 4a, b).

**Fig 4.**
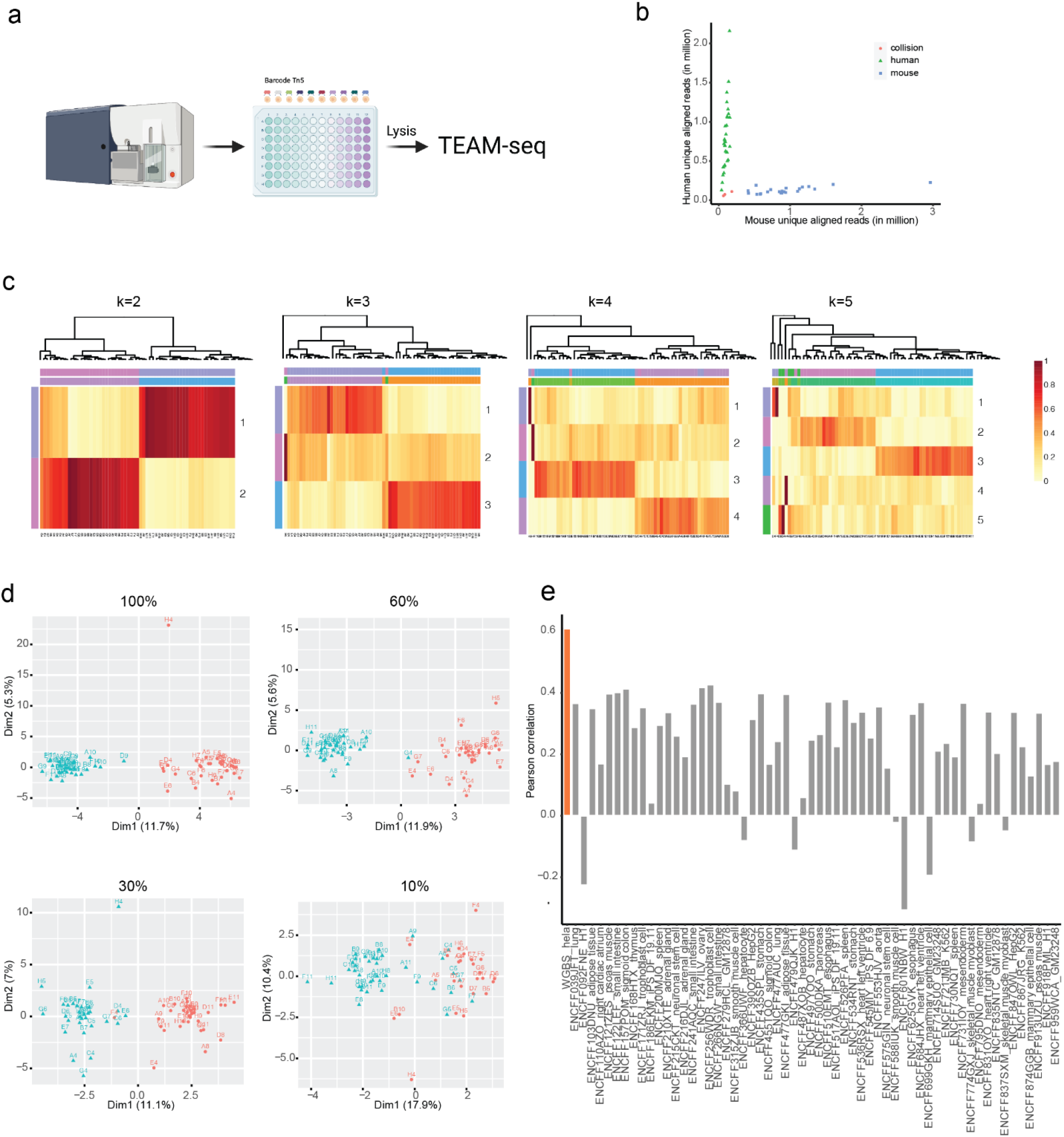
The processes and clustering results of the TEAM-seq on indexed single cells. The cells/nucleuses were sorted into the 96/384-well plate. Then the cells/nucleuses were lysed and tagmentate with indexed transposons. The barcoded genomic fragments were pooled and processed into TEAM-seq. The experimental process and bioinformatic process were described in Supplementalmaterial1_detailedprotocol(a). The barcode collision experiment used 1:1 mixed 4T1/Hela cells and sampled 52 cells (corresponding 52 barcodes)(b). The two barcodes, which did not pass the quality filter (Depth<0.2x), can be classified into neither mouse nor human cells (red). Other barcodes could be classified into either mouse or human cells (1:1 ratio), indicating the barcode collision rate <4%. The cell clustering experiment used 1:1 mixed HEK293T/Hela cells and sampled 52 cells (c,d). The results were summarized as the matrix, including cells (rows) and corresponding methylation ratio in ensemble builds (columns). By the algorithm Non-negative matrix factorization (NMF) ^6^, the matrix V is factorized into (usually) two matrices W and H, with the property that all three matrices have no negative elements (reduce dimension to 2 dimensions). We then studied the most suitable cluster number (k value) for cell classification. As expected, the (k=2) two clusters are propriate for classifying HEK293T and Hela cells. The increasing cluster number did not significantly improve the classification(c). We then down sampled the data from 100% to 10%(d). The two separated cell clusters gradually merged with the decreasing genome coverage, suggesting the importance of the high genome coverage. Also the cell clusters showed the highest correlation with the corresponding cell types from the database.

Next, we profiled an artificial mixture of HeLa cells and the HEK293T cells. We could debarcode >70% reads and **64 cells** (100% cell capturing efficiency), all of which passed the quality filters (depth > 0.2×, mapping rate > **45%**, genome coverage > **3%**). Each cell had a median of 10 million reads and a mean alignment rate of **56%**, approaching the levels seen in the TEAM-seq experiment with 20-pg DNA input (Sfig. 6a). These data translate to the coverage of mappable CpG ranging from ∼5% to ∼20% (Sfig. 6c) and genome coverage ranging from ∼10% to ∼30% (Sfig. 6a). As expected, the sequencing depth was not yet saturated, and an increase in the amount of data could further improve the genome coverage (Sfig. 6b). Among these covered CpG and genome sites, ∼0.2 million Ensembl builds were covered by at least half of the cell population (Sfig. 6f). We did not observe any abnormal coverage bias on the whole genome scale or in CGI (Sfig. 6e, d). Next, we summarized the methylation status of each cell across Ensembl builds to observe the accuracy and variance of methylation mark detection. The correlations between single cells ranged from **0.4 to 0.7** in 0.1–2 kb fragments of Ensembl build (Sfig. 7a). The variance across the genome comprised **0.01–0.05 in CGIs, enhancers, and gene bodies** (Sfig. 7b). On H3K27me3/H3K27ac/H3K4me3, all cells produced the expected nucleosome pattern with corresponding heterochromatin/euchromatin properties (Sfig. 7c).

Next, we profiled a mixture of HEK293T and HeLa human cell lines (Fig. 4). We performed non-negative matrix factorization (NMF) followed by k-means embedding to project cells in two-dimensional space, producing two clearly defined clusters (Fig. 4c, d). We correlated the methylation rate of the collapsed cluster with publicly available WGBS datasets (Fig. 4e). One cell cluster showed the highest correlation with the corresponding cell type (Fig. 4e). To display the advantage of increased coverage of the single cell, we down-sampled the coverage and found that the identified two cluster distances decreased with coverage until merging (Fig. 4d). Overall, TEAM-seq could generate high methylation site coverage on the indexed single cells in a high-throughput manner.

## Discussion

In the current study, we developed the TEAM-seq method based on the enzymatic cytidine conversion (EM-seq) and T7 polymerase amplification, achieving a very high genome coverage in 100 pg and 20 pg input samples, and even in single cells. However, EM-seq still demonstrated 0.7% lower conversion rates than the gold standard BS-seq, resulting in a higher false-positive/negative rate. The higher number of false positive/negative methylation marks may be caused by the substrate/motif preference of Tet1/2 and APOEC3A or different enzymatic efficiency. The slightly higher false positive/negative rate would be problematic in single-cell analysis. Because a single cell has only one/two copies of genome, the accuracy is difficult to correct. However, this problem cannot be easily avoided in enzymatic conversion. Given the significant DNA loss during the harsh bisulfite treatment, we believe that enzymatic conversion is a better choice for single-cell methylation analysis. Moreover, we are developing a machine learning algorithm that may help distinguish false positive/negative methylation marks in single cells.

We have tried many other approaches in the current study, but most methods produced surprisingly unsatisfactory results. The Trueprime kit demonstrated superior performance in single-cell genome amplification compared to that achieved with MDA, but not for enzymatically converted uridine-rich DNA. We suspected that heavy modifications (5CaC/5fC/5gmC) generated by EM-seq inhibited primer synthesis by primase. A similar phenomenon was also observed during the GpC methyltransferase treatment, which preserved 25% of the GC content in the enzymatic conversion. The high GC content adversely affected MDA owing to the parallel increase in heavy modifications (5CaC/5fC/5gmC). We also observed a related phenomenon in a mixture of HEK293T and HeLa cells. HEK293T cells generated twice more reads than HeLa cells, suggesting the preferential amplification of HEK293T cell DNA. This bias was not observed in a mixture of HeLa and 4T1 cells. Considering the higher abundance of 5mC marks in cancer cells^25^, heavy modifications may also lead to the inefficiency of T7 polymerase. Therefore, the vast global difference in the methylation rate may cause biased amplification in cell populations for pooling the indexed cells.

In TEAM-seq, we used Tn5 to fragment genomic DNA, but the fragment length was difficult to control precisely in single cells. The varied fragment length required nanopore or Pacbio sequencing. The nanopore method has higher error rate than Pacbio. However, we found that two reads per site could confidently correct the sequencing errors. Otherwise, our patented linked-reads method ^26^ on the Pacbio platform could be an alternative to nanopore sequencing, which could increase 5x output of Pacbio sequencing.

Overall, TEAM-seq reported genome coverage for ultralow input DNA amounts and DNA from single cells. This method could help in studies of samples with a limited DNA amount, e.g., embryos, oocytes, clinical tissues, etc. The high coverage of single cell genomes may further disclose methylation heterogeneity of different cell populations.

## Methods

### Experimental methods

#### Assess the EM-seq and WGBS

*Arabidopsis* DNAs were given by the other lab. The EM-seq was performed and NEB EM-seq protocol (NEB E7120S)(1ug DNA per reaction). The WGBS was parallelly performed by using MGIEasy Whole Genome Bisulfite Sequencing Library Prep Kit (MGI 1000005251). Both methods were similar except the conversion steps. The final products were sequenced by MGISEQ-2000.

#### MDA for the converted DNA

Both of the WGBS and EM-seq converted cytidine for the constructed DNA library, which was fragmented and ligated with adaptors. Unlike the WGBS, MDA-EM-seq used the Phi29 to do the multiple displacement amplification on the non-fragmented DNA. The genomic DNA were directly processed to EM-seq conversion steps (https://international.neb.com/protocols/2019/03/28/protocol-for-use-with-standard-insert-libraries-370-420-bp-e7120, 1.5-1.9). The converted single-strand DNAs were around 6∼10kb after enzyme treatment. The clean-up single-strand DNAs were processed to the MDA amplification by REPLI-g Single Cell Kit (Qiagen 150343). For Trueprime MDA-EM-seq, the clean-up DNAs underwent the protocol SYGNIS TruePrime^TM^ WGA & Single Cell WGA Kits (Lucigen), instead of the REPLI-g single cell kit. In the EM-MDA-seq with additives, all the addictive (3% PEG,100mM TMAC, 3%PEG+100mM TMAC) were added to the REPLI-g Single Cell reaction. In the EM-MDA-seq with BST amplification, we used the BST2.0 (NEB M0537S) to substitute the Phi29 in REPLI-g single cell kit. The amplified DNAs were processed as MGIEasy Universal DNA Library Prep Kit V1.0 and sequenced in MGISEQ-2000.

#### scWGBS_ng

The scWGBS_ng was downloaded ^27^. Briefly, bisulfite conversion was performed using the EZ DNA Methylation-Direct Kit (Zymo Research, D5020) according to the manufacturer’s protocol, with the modification of eluting the DNA and minimizing the DNA loss. The single stranded library construction was followed.

#### TEAM-seq

We first fragment the DNA by Tn5 transposon with biotin adaptor. After tagmentation, the DNA with biotin adaptors could be bound with the streptavidin beads. Then the DNAs carried on streptavidin beads were processed to EM-seq (https://international.neb.com/protocols/2019/03/28/protocol-for-use-with-standard-insert-libraries-370-420-bp-e7120, 1.5-1.9). The DNA immobilized on the beads could be smoothly transfer to the next reaction without the significant loss. After conversion, the single strand DNAs (beads-on reaction) were extended on the complementary oligos to generate the T7 promoters. Then we performed the bead-on T7 transcription reaction by HiScribe™ T7 High Yield RNA Synthesis Kit (NEB E2040S). The amplified RNAs were processed to the PCR-cDNA sequencing kit (nanopore). The detailed description of methods can be found in Supplemental Material supplementalmaterial1_detailedprotocol. For the single cell TEAM-seq, we used the Qiagen protease to lysis the single cell to release DNAs and heat inactivated the Qiagen protease before processing to the Tn5 tagmentation.

#### Cell culture

Human mammary gland carcinoma cell line HeLa/HEK293T were obtained from ATCC. MCF-7 were grown in DMEM (Gibco,11995065) supplemented with 10% FBS (Gibco,10099141), 0.01mg/ml insulin (HY-P1156, MedChemExpress), and 1% penicillin-streptomysin (Gibco, 15140122). Cell line was regularly checked for mycoplasma infection (Yeasen, 40612ES25).

### Bioinformatics method

#### Data processing

Sequencing adapters were cut using fastp (v0.21.0). The remaining reads were trimmed based on their theoretical barcode locations. Then they were aligned to the barcode white list in the following pattern: Ns-GGGAGATACAACCTACAATCACT-10bp barcode- AAATATATATAAAAAACAA-Ns using bowtie2 local alignment mode (v2.3.4). Only reads aligned to the barcode reference with high mapping quality would be processed and used for subsequent analysis.

To assess the accuracy of demultiplexing, 10,048 reads were simulated with a length of 52 bases, including a 23bp 5’ primer, 10bp barcode from the white list, and a 19bp 3’ primer. In order to intimate nanopore sequencing errors, a mean indel rate of 2.96% and a mean mismatch rate of 2.77% were introduced to the simulated data.

Reads were aligned by methylpy (v1.3) in the following steps: (1) reads mapped to the forward strand GRCh37 reference genome were de-duplicated with the parameter “--pbat --remove-clonal --path-to --picard --min-mapq 15”. (2) Remaining unmapped reads were then mapped to the reverse strand of the reference genome in the same way. (3) Merge all mapped bam, then call methylation status by methylpy call_methylated_sites. In terms of human-mouse mixed library, a combined human-mouse hybrid genome was used instead of GRCh37 reference.

Fastq files of scWGBS, sci-MET, snmC-seq and scBS-seq downloaded from the database were aligned to the reference genome and called the methylation ratio using Bismark (v0.22.3). Any other conventional WGBS data was analysed using our standard pipeline.

#### Analysis of cytosine modifications called by TEAM-seq, scWGBS and WGBS

Chromosomal methylation status was plotted using CG/CHG/CHH positions covered by more than three times. The chromosome was then binned into equal sizes (100kb for Arabidopsis Chr4, 2Mb for human Chr1 and Chr4). Mean methylation ratio per bin was calculated and plotted along the chromosome. The mean occurrences of C/CG/CHG/CHH in each bin was computed and plotted in the same way.

Information of histone (H3K27me3/H3K27ac/H3K4me3) ChIP-seq peaks was downloaded from the ENCODE database. Bed files of gene elements including CGIs, TSS, TES and gene body were downloaded from UCSC genome browser. These regions were binned into 20 or 50 windows, flanking by 50 windows of the up-/downstream regions. The methylation ratio of each bin was averaged and plotted using R.

#### Average coverage depth in CGI

Total nucleotide count for each bin was reported using samtools bedcov (v1.9), then average coverage depth in each window of the CGI region was computed for TEAM-seq, scWGBS and WGBS. To overcome differences in sequencing depth, TEAM-seq and WGBS data were down-sampled to the corresponding sequencing depth of scWGBS. Average coverage depth for single cell data was computed in the same manner.

#### Correlation and consistency analysis

For each single cell, weighted methylation ratios of the Ensembl Regulatory Build regions were computed in a previously described method ^28 10^. We then calculated the Pearson correlation of regions including all Ensembl Regulatory loci, promoter region, enhancer region and CTCF binding sites. Clustered heatmap was then plotted using pheatmap in R.

In order to correlate with bulk WGBS data, we merged the methylation information of all cells in cluster 1 and calculated methylation ratio at single nucleotide level. The methylation ratio was then correlated to WGBS datasets available from the ENCODE database as well as the conventional HeLa WGBS data ^29^.

The consistency of methylation status was compared between WGBS and EM-seq, TEAM-seq 100pg and 1ug WGBS, TEAM-seq 20pg and WGBS, and among three TEAM-seq 20pg libraries. For each comparison, only positions covered more than three times were used and a methylation ratio of 0.5 was used as the threshold. For example, if the methylation ratio of a cytosine is either below 0.5 or above 0.5 in both libraries, the methylation pattern was considered as consistent. Venn diagram and Euler diagram were plotted using VennDiagram and eulerr packages in R.

#### NMF decomposition and k-means clustering

Non-negative Matrix Factorization (NMF) is a decomposition algorithm to split a matrix V into two matrices W (dictionary matrix) and H (meta-feature matrix). Due to its clustering property, it has been widely used in DNA methylation studies to identify features that contribute the most to the cluster. Therefore, using Ensembl Regulatory Build as the feature, NMF clustering was performed to distinguish HeLa and HEK293T cell lines. Then, k-means clustering algorithm was used to cluster cells by selecting 100%, 60%, 30% and 10% features to investigate the trend of two clusters getting more similar to each other. All clustering analyses were performed in R.

## Data availability

TEAM-seq_100pg, TEAM-seq_20pg, HeLa&4T1 mixture data as well as HeLa&HEK293T mixture data are available at China National GeneBank (CNGB) ^30, 31^ with project number of CNP0002270.

## Supporting information

Supplementalmaterial1_detailedprotocol1

## Acknowledgment

This research was supported by the Science, Technology, and Innovation Commission of Shenzhen Municipality (grant number JSGG20170824152728492). The supporter had no role in designing the study, data collection, analysis, interpretation, or writing of the manuscript.

## Author contributions

CT designed and supervised the experiments. JW performed the experiments; YTF, FC, and CT performed bioinformatics data analysis. All authors collectively analyzed experimental data. All authors read and approved the final draft of the manuscript.

---

**Sfig1.**
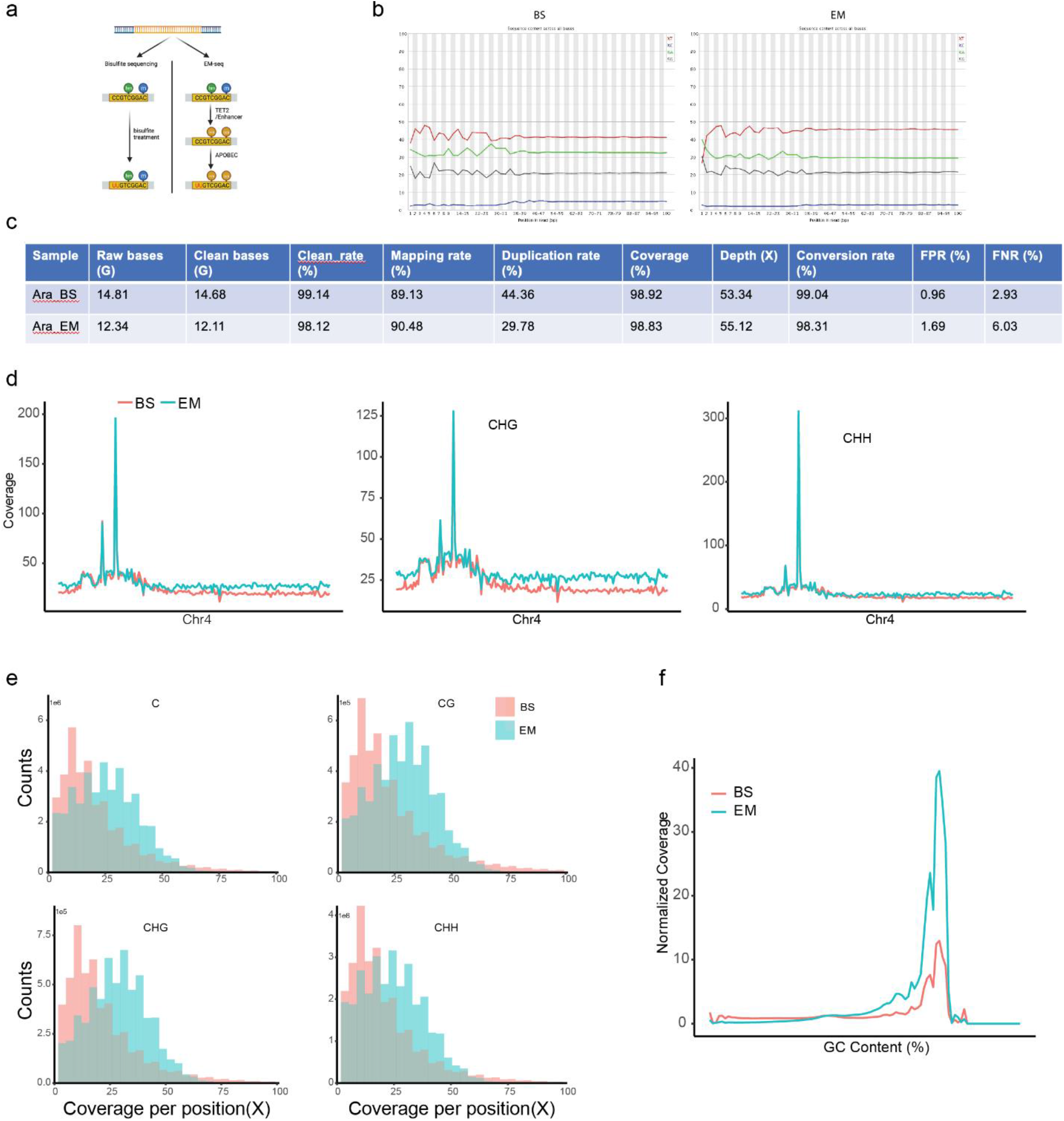
The quality control of the EM-seq and BS-seq. We showed the schematic plot of the mechanism of EM-seq and BS-seq (a). Bisulfite treatment could convert all the unmethylated cytidine(C) to uridine(U) without affecting methylated cytidine (5mC). The 5mC could be preserved and readed as “C” in the sequencing. EM-seq used the Tet2/enhancer to transform the 5mC/5hmC to 5caC,5fC, and 5gmC, which could be preserved in the APOEC3A treatment. APOEC3A then deaminate the cytidine to uridine, and cannot convert 5caC,5fC,5gmC to uridine. The preserved “5mC” signal could also be recognized as “C” in sequencing. The phenotype of these two types of sequencing data was similar, so the analysis pipelines are the same. In the sequencing data, both EM-seq and BS-seq demonstrated the similar sequencing base bias (b). The sequencing quality and data yield are comparable between EM-seq and BS-seq. The final clean data yield was similar at 98%. EM-seq mapping rate was slightly higher. Duplication rate in EM-seq is 50% fewer than in the BS-seq. The global genome coverages were similar between these two methods at 98%. The unmodified cytidine conversion rates were 99.04% in BS-seq and 98.31% in EM-seq. We used the negative control unmodified lambda DNA to calculate the false positive rate (FPR) and the positive control synthetic DNA with specific modified site to calculate the false negative rate (FNR). FPR=C in negative lambda DNA/(C+U in negative lambda DNA);FNR=U in positive PTXB1/(C+U in positive PTXB1) (c). We further look the coverage distribution on Chr4. In the whole genome view, the EM-seq coverage depth are generally higher than the BS-seq in all typical methylation pattern (d). The EM-seq could cover C/CG/CHH/CHG around 25x medium coverage comparing with BS-seq 12x medium coverage(e). To further analyze the relationship between the coverage and GC content. We found both EM-seq and BS-seq tend to cover the areas with medium GC content (50%∼75%) (f).

**Sfig2.**
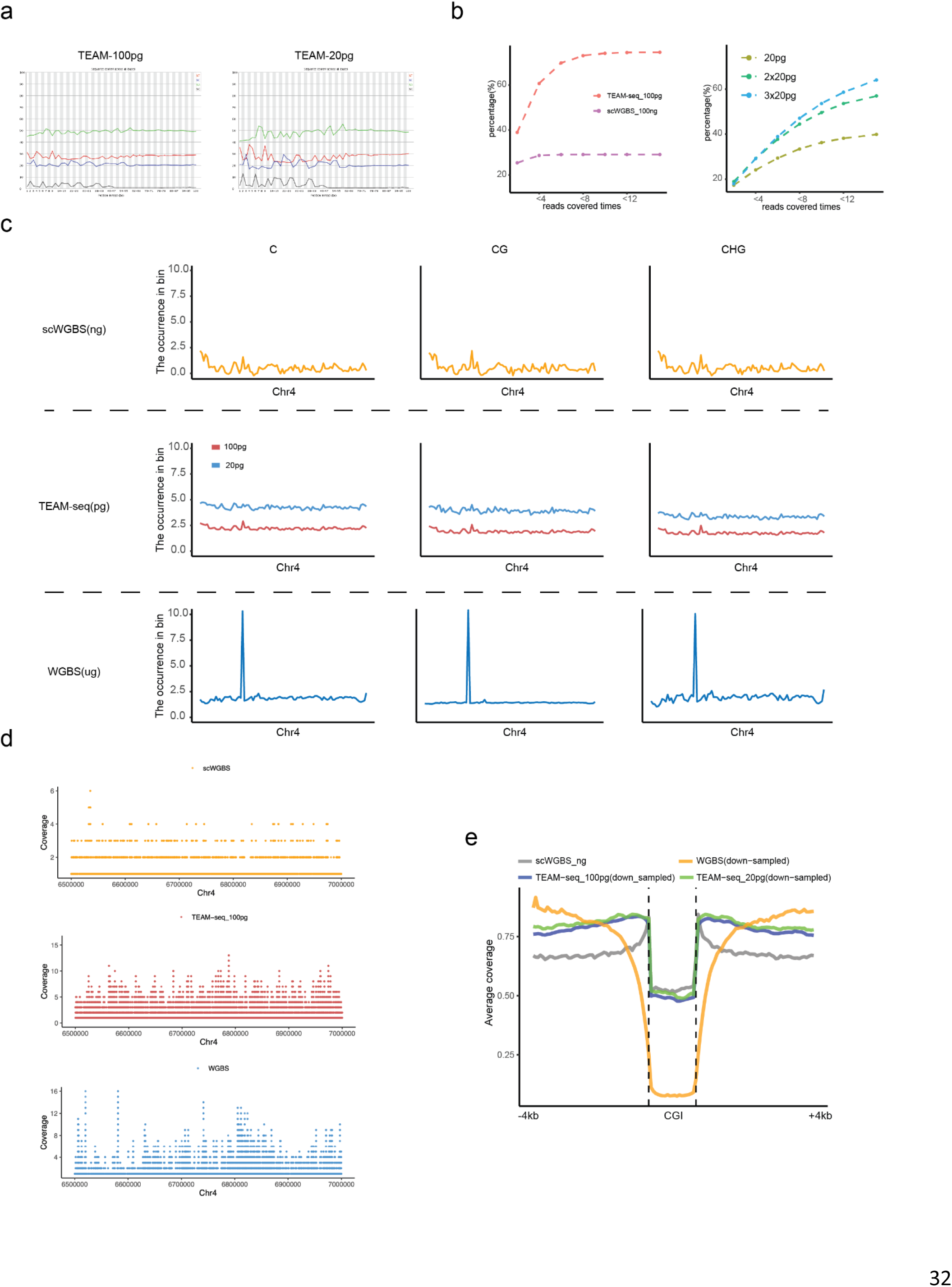
The sequencing quality and the even coverage of the ultra-low input TEAM-seq.(a) The lines with four colors showed the ATGC proportion in each sequencing base. The TEAM-seq_100pg and TEAM-seq_20pg represented the sequencing result with different initial DNA input 100-pg/20-pg. The 20-pg TEAM-seq showed the more severe fluctuation than the 100-pg TEAM-seq. (b) Left panel showed the accumulation curve of the read covered times per base in 100-pg TEAM-seq and 100-ng scWGBS. The scWGBS used 100-ng as initial DNA input and simplified the experiments to one tube reaction to minimize the loss of operation, and used single stranded library construction ^7^. In the TEAM-seq_100pg, 70% sites were covered fewer than 7 times and outperformed the scWGBS_100ng. The scWGBS get to the saturation point (30% genome sites) at 4. We used 20pg genomic DNAs, as the ultra-low input, close to the single cell level. These experiments were repeated for three times. Then we merged these sequencing results of the three 20-pg experimental repeats. The accumulation curve got greatly improvement after merging two 20-pg experimental repeats. (c) The occurrences of covered C/CG/CHG (100 bins) on the chromosome 4 among scWGBS, TEAM-seq, WGBS. The WGBS used 1ug genomic DNA as initial input. All four data were down sampled to the same level. The scWGBS_100ng showed the uneven genomic coverage and some areas were not covered. The WGBS_1ug had the even coverage on the genome with the serious coverage bias on the specific region. Comparing with these two methods, TEAM-seq demonstrated the very even coverage on the Chr4, with ultra-low input DNA. (d) The coverage on the region Chr4:6500000-7000000. TEAM-seq demonstrated the even coverage on this region. (e) The average coverage around the CGI region among scWGBS_100ng, WGBS_1ug, TEAM-seq_100pg, TEAM-seq_20pg. The data were down sampled to the same sequencing depth. The TEAM-seq_20pg demonstrated the highest genome coverage among these three methods.

**Sfig3.**
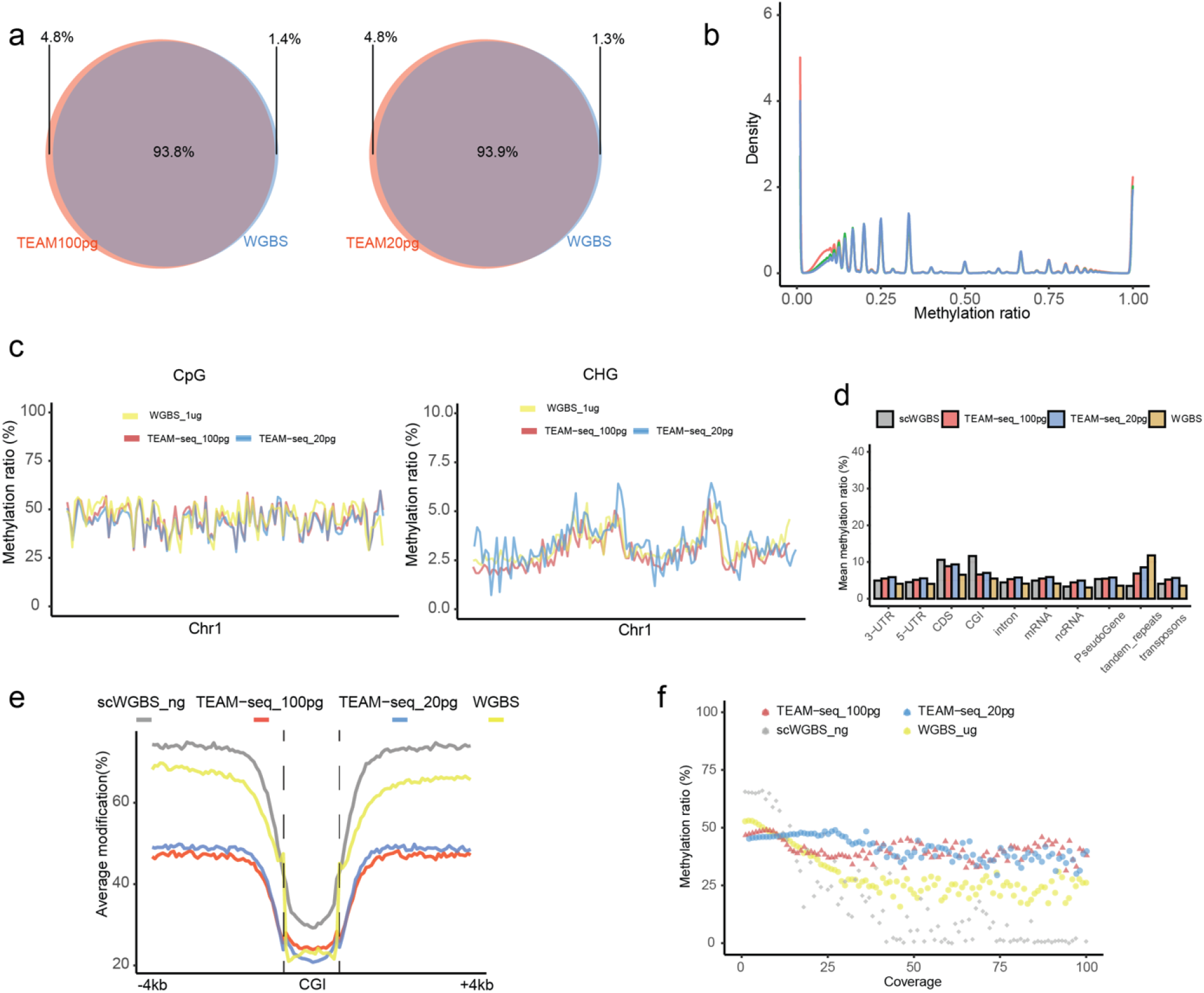
The TEAM-seq and WGBS showed the high consistency in the methylation detection. We selected the high coverage sites (covered by more than 3 reads). The sites with >50% methylation ratio counted as the methylated sites and the sites with <50% methylation ratio counted as the unmethylated sites. The methylation status consistency (methylated/unmethylated) was over 93% between TEAM-seq (20-pg/100-pg) and WGBS (1ug). The calculated methylation ratio correlation between 100-pg TEAM-seq and WGBS was around 0.82 (a). Due to the 20pg genome DNA only contain 1-5 copies of the genomic segments, the methylation ratio was at both of the extremes (0,1) (b). The over methylation level distributions on chr1 (CpG, CHG) were similar between TEAM-seq(20-pg/100-pg) and WGBS(1ug) (c). The TEAM-seq average methylation ratios on the gene element were slightly higher than the WGBS, expect the tandem_repeats (d). We used the CGI as example to study the detailed methylation distribution on the CGI. These four methods all showed the depressed methylation on the CGI and high methylation on the CGI shoulders. The TEAM-seq display 30% lower methylation on the CGI shoulders (e). The scWGBS demonstrated the most serious methylation bias resulted by the coverage (f). In the contrast, the TEAM-seq both 100pg and 20pg have the very slightly coverage related methylation bias. The results were similar to the previous report TAPS ^8^.

**Sfig4.**
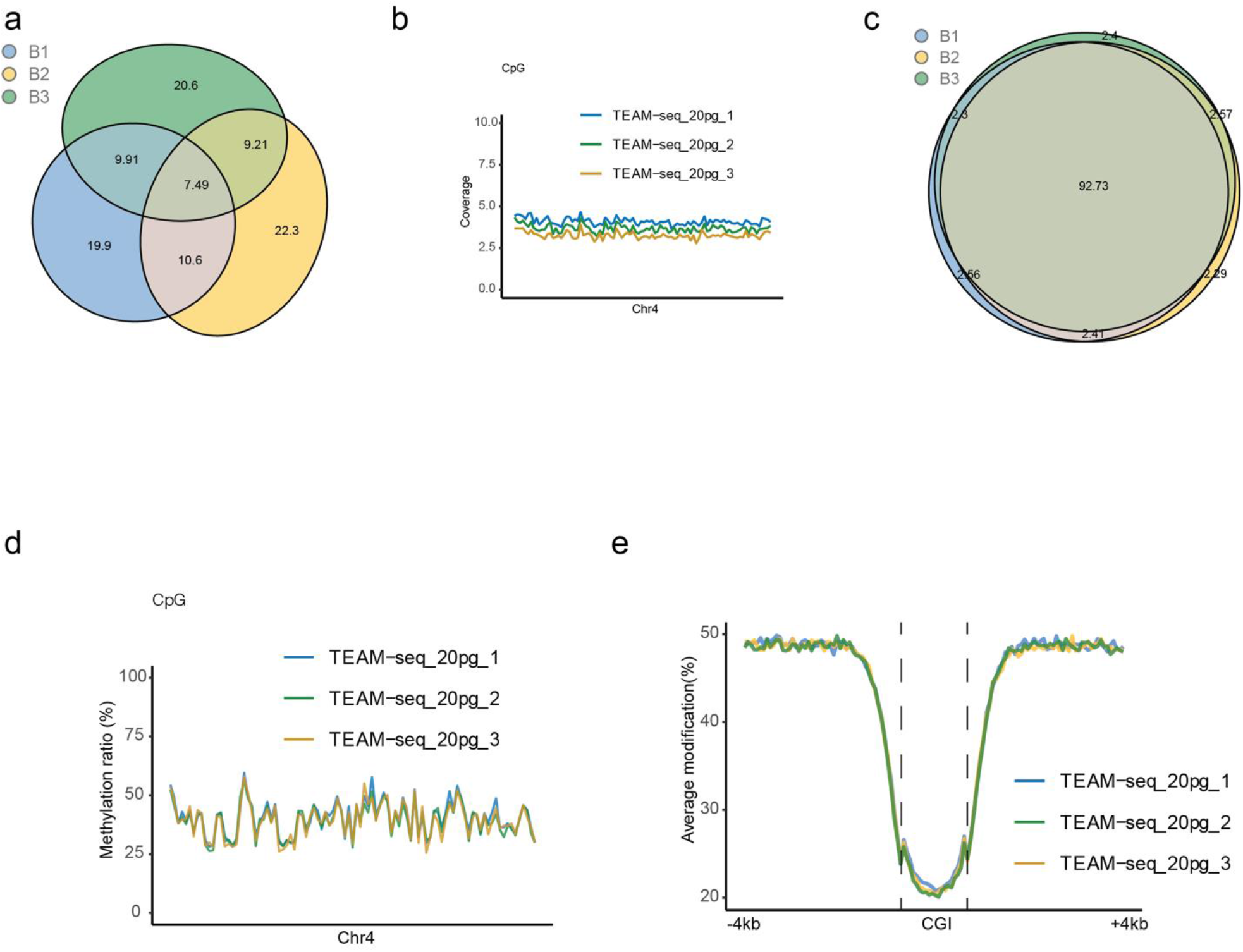
The high reproducibility in ultra-low input TEAM-seq (20-pg). The 20-pg TEAM-seq was repeated three times independently. The common genome sites, shared by these three independent samples, were 7.49%, due to the sampling bias with minimal amount DNAs (a). The three replicates displayed the similar genome coverage on chr4 (b). The methylation statuses were very consistent between these three replicates (consistency definition as Fig.3). Due to the high methylation consistency, the CpG methylation ratio distribution of these three replicates showed the same pattern on chr4(d). Especially on the CGIs(e), the methylation ratio distributions were nearly the same among these three replicates.

**Sfig5.**
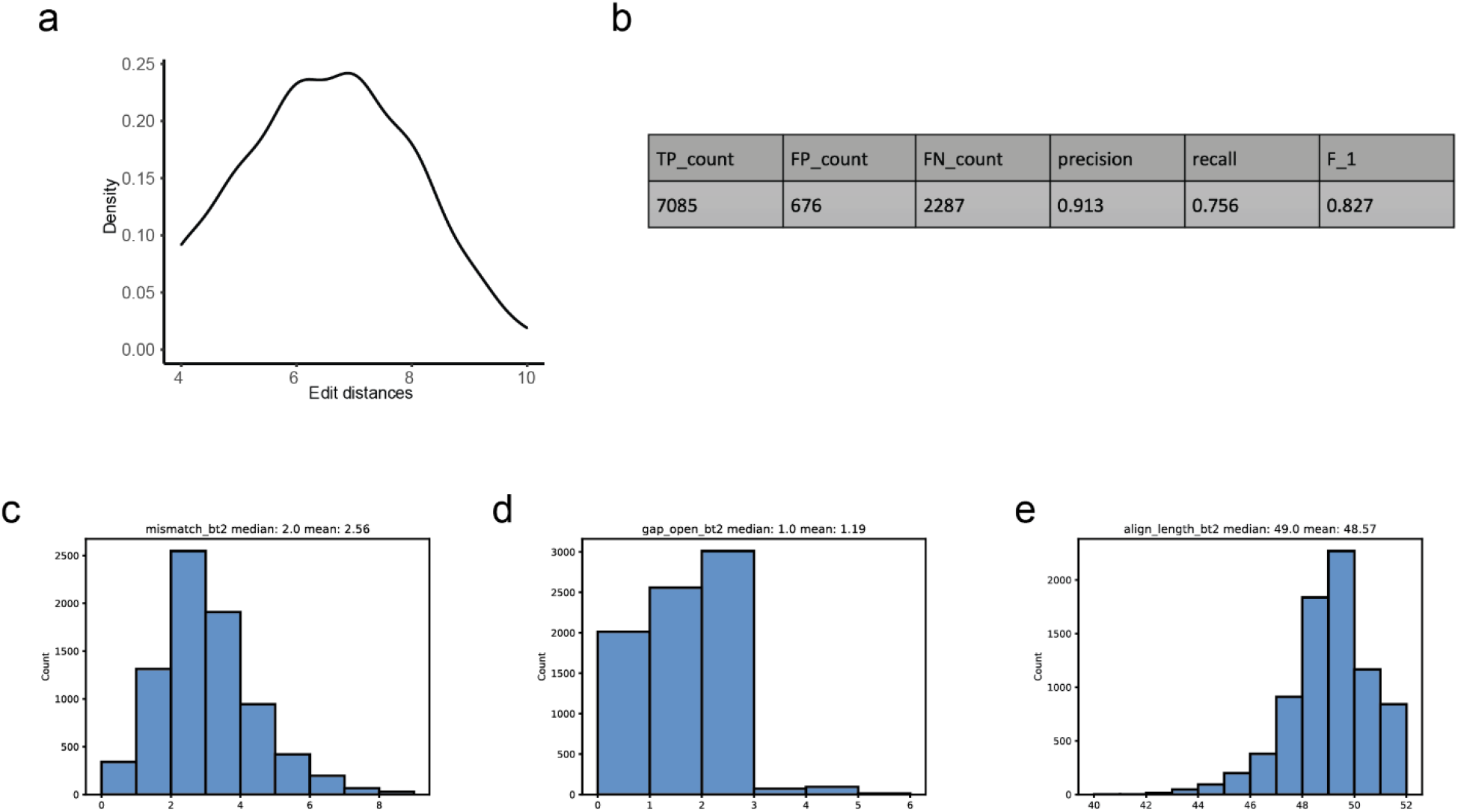
The barcode designed distance and assessment of the debarcoding methods. We designed the 10 nucleotide barcodes, which own the 4∼10 letter differences between each other. The medium edit distance is round 7 (Hamming distance)(a). Then we simulated the nanopore synthetic data, to create enough mismatches, insertions, deletions in barcode and adaptor sequences. We used our house pipeline (method) to debarcode the synthetic data and calculated the TP(True positive), FP(False positive), FN(False negative), precision, recall and F1 (b). To compare the simulated sequence with its original templates, the simulation generated the mean number of mismatches 2.56(c) and the mean number of gaps 1.19(d) on the barcode and adaptor sequence (e, the mean of barcode plus adaptor length 48.57).

**Sfig6.**
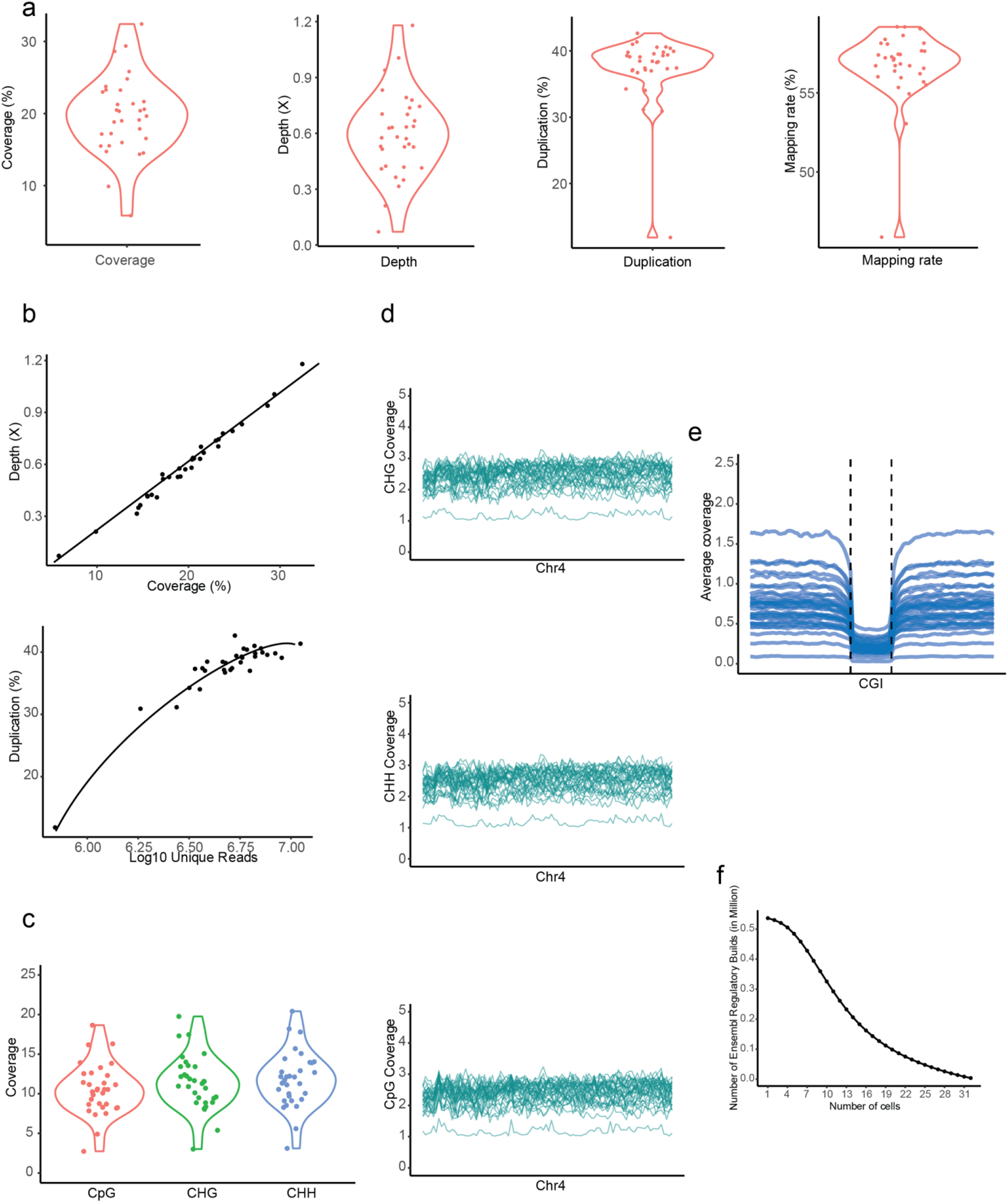
The coverage of the TEAM-seq on the indexed single cells (HEK293T cells). The medium genome coverage is around ∼20% on average ∼0.6X sequencing depth. The average duplication rate and mapping rate were ∼38% and ∼60%(a) (The coverage/depth calculations were described on method). The sequencing depth has the significant effect on the genome coverage with the linear increasing trend, suggesting that the sequencing depth is not close to saturation. The more sequencing data could further improve the genome coverage(b). The duplication rate is hitting the plateau ∼38% with 0.6x depth (10^7^ reads) (b). The mean CpG/CHG/CHH coverage is around ∼12%/13%/13%(c). On the whole genome scale, each cell showed the uniform coverage on chr4, which is like 100-pg/20-pg TEAM-seq(d). Moreover, each cell showed the similar coverage pattern on CGI with various sequencing depth(e). Most of the single cell algorithm rely on the common areas covered by most cells, which are important in single cell analysis. In scTEAM-seq (single cell TEAM-seq), ∼0.2million Ensembl builds could be covered half cellular population(f).

**Sfig7.**
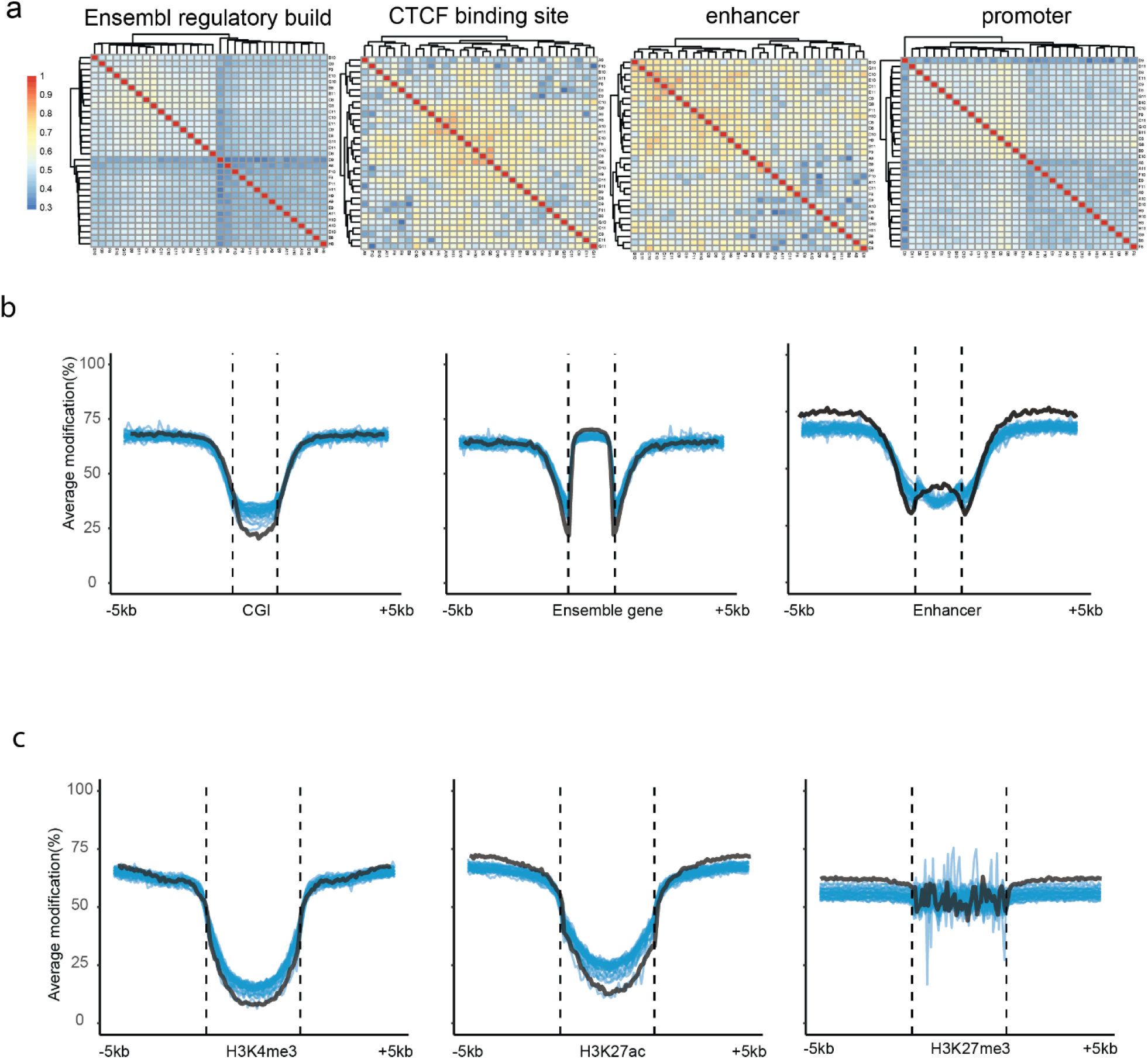
The accuracy and consistency of the TEAM-seq on the indexed single cells (HEK293T). The single cell methylation is summarized as the matrix, including the cells (row) and the mean methylation of ensemble builds. The Ensembl builds carried the location information of CpG islands, enhancers, promoters, CTCF binding motifs. We then calculated the Pearson correlations between each cell among these builds. Half of the single cells demonstrated high correlations in the CTCF binding motifs (∼0.7) and enhancers (∼0.78) (a). Then we plotted the methylation distribution pattern of each cell (each blue line represented one cell) on gene elements(b), including CGIs/gene bodies/enhancers, and histone modification motifs(c), including H3K4me3(active)/H3K27ac(active)/H3K27me3(repressive). The black line indicated the methylation distribution of scWGBS (100ng). The methylation distributions were similar between scWGBS and indexed single cell TEAM-seq.

## Bibliography

1. Hwang, B., Lee, J.H. & Bang, D. Single-cell RNA sequencing technologies and bioinformaticspipelines. Experimental & Molecular Medicine 50, 1–14 (2018).

2. Nawy, T. Single-cell sequencing. Nature Methods 11, 18–18 (2014).

3. Li, E. Chromatin modification and epigenetic reprogramming in mammalian development. Nat Rev Genet 3, 662–673 (2002).

4. Bird, A. Perceptions of epigenetics. Nature 447, 396–398 (2007).

5. Tollefsbol, T.O. Methods of epigenetic analysis. Methods Mol Biol 287, 1–8 (2004).

6. Guo, H. et al. Profiling DNA methylome landscapes of mammalian cells with single-cell reduced-representation bisulfite sequencing. Nat Protoc 10, 645–659 (2015).

7. Guo, H. et al. Single-cell methylome landscapes of mouse embryonic stem cells and early embryos analyzed using reduced representation bisulfite sequencing. Genome Res 23, 2126–2135 (2013).

8. Karemaker, I.D. & Vermeulen, M. Single-Cell DNA Methylation Profiling: Technologies and Biological Applications. Trends in Biotechnology 36, 952–965 (2018).

9. Miura, F., Enomoto, Y., Dairiki, R. & Ito, T. Amplification-free whole-genome bisulfite sequencing by post-bisulfite adaptor tagging. Nucleic Acids Research 40, e136–e136 (2012).

10. Smallwood, S.A. et al. Single-cell genome-wide bisulfite sequencing for assessing epigenetic heterogeneity. Nature Methods 11, 817–820 (2014).

11. Miura, F. & Ito, T. Highly sensitive targeted methylome sequencing by post-bisulfite adaptor tagging. DNA Research 22, 13–18 (2015).

12. Kobayashi, H. et al. Repetitive DNA methylome analysis by small-scale and single-cell shotgun bisulfite sequencing. Genes to Cells 21, 1209–1222 (2016).

13. Farlik, M. et al. Single-Cell DNA Methylome Sequencing and Bioinformatic Inference of Epigenomic Cell-State Dynamics. Cell Reports 10, 1386–1397 (2015).

14. Luo, C. et al. Single-cell methylomes identify neuronal subtypes and regulatory elements in mammalian cortex. Science 357, 600–604 (2017).

15. Mulqueen, R.M. et al. Highly scalable generation of DNA methylation profiles in single cells. Nat Biotechnol 36, 428–431 (2018).

16. Liu, Y. et al. Bisulfite-free direct detection of 5-methylcytosine and 5-hydroxymethylcytosine at base resolution. Nature Biotechnology 37, 424–429 (2019).

17. Frommer, M. et al. A genomic sequencing protocol that yields a positive display of 5-methylcytosine residues in individual DNA strands. Proc Natl Acad Sci U S A 89, 1827–1831 (1992).

18. Vaisvila, R., et al. EM-seq: Detection of DNA Methylation at Single Base Resolution from Picograms of DNA. bioRxiv, 2019.2012.2020.884692 (2020).

19. Vaisvila, R. et al. Enzymatic methyl sequencing detects DNA methylation at single-base resolution from picograms of DNA. Genome Res 31, 1280–1289 (2021).

20. Dean, F.B., Nelson, J.R., Giesler, T.L. & Lasken, R.S. Rapid amplification of plasmid and phage DNA using Phi 29 DNA polymerase and multiply-primed rolling circle amplification. Genome Res 11, 1095–1099 (2001).

21. Huang, L., Ma, F., Chapman, A., Lu, S. & Xie, X.S. Single-Cell Whole-Genome Amplification and Sequencing: Methodology and Applications. Annu Rev Genomics Hum Genet 16, 79–102 (2015).

22. Picher, Á.J. et al. TruePrime is a novel method for whole-genome amplification from single cells based on TthPrimPol. Nature Communications 7, 13296 (2016).

23. Choi, J.Y., Lim, S., Eoff, R.L. & Guengerich, F.P. Kinetic analysis of base-pairing preference for nucleotide incorporation opposite template pyrimidines by human DNA polymerase iota. J Mol Biol 389, 264–274 (2009).

24. Chen, C. et al. Single-cell whole-genome analyses by Linear Amplification via Transposon Insertion (LIANTI). Science 356, 189–194 (2017).

25. Ehrlich, M. DNA methylation in cancer: too much, but also too little. Oncogene 21, 5400–5413 (2002).

26. Zheng, Y.-F., et al. HIT-scISOseq: High-throughput and High-accuracy Single-cell Full-length Isoform Sequencing for Corneal Epithelium. bioRxiv, 2020.2007.2027.222349 (2020).

27. Farlik, M. et al. Single-cell DNA methylome sequencing and bioinformatic inference of epigenomic cell-state dynamics. Cell Rep 10, 1386–1397 (2015).

28. Zerbino, D.R., Wilder, S.P., Johnson, N., Juettemann, T. & Flicek, P.R. The ensembl regulatory build. Genome Biol 16, 56 (2015).

29. An integrated encyclopedia of DNA elements in the human genome. Nature 489, 57–74 (2012).

30. Chen, F.Z. et al. CNGBdb: China National GeneBank DataBase. Yi Chuan 42, 799–809 (2020).

31. Guo, X. et al. CNSA: a data repository for archiving omics data. Database (Oxford) 2020 (2020).

## References

1. Smallwood, S.A. et al. Single-cell genome-wide bisulfite sequencing for assessing epigenetic heterogeneity. Nature Methods 11, 817–820 (2014).

2. Miura, F. & Ito, T. Highly sensitive targeted methylome sequencing by post-bisulfite adaptor tagging. DNA Research 22, 13–18 (2015).

3. Kobayashi, H. et al. Repetitive DNA methylome analysis by small-scale and single-cell shotgun bisulfite sequencing. Genes to Cells 21, 1209–1222 (2016).

4. Farlik, M. et al. Single-Cell DNA Methylome Sequencing and Bioinformatic Inference of Epigenomic Cell-State Dynamics. Cell Reports 10, 1386–1397 (2015).

5. Luo, C. et al. Single-cell methylomes identify neuronal subtypes and regulatory elements in mammalian cortex. Science 357, 600–604 (2017).

6. Mulqueen, R.M. et al. Highly scalable generation of DNA methylation profiles in single cells. Nat Biotechnol 36, 428–431 (2018).

7. Farlik, M. et al. Single-cell DNA methylome sequencing and bioinformatic inference of epigenomic cell-state dynamics. Cell Rep 10, 1386–1397 (2015).

8. Liu, Y. et al. Bisulfite-free direct detection of 5-methylcytosine and 5-hydroxymethylcytosine at base resolution. Nature Biotechnology 37, 424–429 (2019).

