## Supplementalmaterial1_detailedprotocol1 for "High coverage of single cell genomes by T7-assisted enzymatic methyl-sequencing": Supplementalmaterial1_detailedprotocol 1.docx

### A. Primer sequence

| Primer name | Primer sequence |  |
| --- | --- | --- |
| Template extension primer(T7 oligo) | CATATAATACGACTCACTATAGGGAGATACAACCTACAATCACT |  |
| RT primer | ACTAGTAAGCAGTGGTATCAACGCAGAGTACAGATGTGTATAAGAGATAG |  |
| Long primerP5 | AATGATACGGCGACCACCGAGATCTACACGGGAGATACAACCTACAATCACT | 5-pho |
| Long primerP7 | CAAGCAGAAGACGGCATACGAGATGAGTACAGATGTGTATAAGAGATAG |  |
| Short P5 | AATGATACGGCGACCACCGAGATC | 5-pho |
| Short P7 | CAAGCAGAAGACGGCATACGAGAT |  |
| New Tn5-pho | CTGTCTCTTATACACATCT | 5-pho |
| New bio-bottom | TACAACCTACAATCACT NNNNNNNNNN AGATGTGTATAAGAGACAG | 5'-Biotin |

### B. The sequence through the whole procedure(TEAM-seq):

1.Tagmentation: (Green-ME sequence for Tn5, Red-adaptor sequence)

TACAACCTACAATCACTTTATTAACAAAGATGTGTATAAGAGACAG	CTGTC TC TTATA CACATCT 3

TCTACACAT ATTC TC TGTC	GACAGAGAATATGTGTAGAAACAATTATTTCACTAACATCCAACAT 5

2. Gap filling:

TACAACCTACAATCA CTTTATTAACAAAGATGTGTATAAGAGACAG CTGTC TC TTATA CACATCTTTGTTAATAAAGTGATTGTAGGTTGTA 3

ATGTTGGATGTTAGTGAAATAATTGTTTCTACACAT ATTCT CTG TC GACAGAGAATATGTGTAGAAACAATTATTTCACTAACATCCAACAT 5

3. EM conversion:

TAUAAUUTAUAATUAUTTTATTAAUAA AGATGTGTATAAGAGAUAG UTGTU TU TTATAUAUATUTTTGTTAATAAAGTGATTGTAGGTTGTA 3

&

ATGTTGGATGTTAGTGA AATAATTGTTTUTAUAUATATTU TUTGTU GAUAGAGAATATGTGTAGAAAUAATTATTTUAUTAAUATUUAAUAT 5

4. Phi29 or Q5 extension on the T7 oligo (Bold-T7 promoter sequence, cyan-the sequence to align)

5‘ T7 oligo

TCACTAACATCCAACATAGA**GGGATATCACTCAGCATAAT**ATAC

TAUAAUUTAUAATUAUT TTATTAAUAAAGATGTGTATAAGAGAUAG	UTGTU TU TTATAUAUATUTTTGTTAATAAAGTGATTGTAGGTTGTA	3‘

After extension：

*ATATTAAATA TTAATAA AATAATTATT TCT ACACAT ATTC TC TA TC* ………………………. *AACAAAAAATATAT A TAAA*ACAATTATTTCACTAACATCC AACAT AGA**GGGATATCACTCAGCATAAT**ATAC 5

TAUAAUUTAUAATUAUTTTATTAAUAAAGATGTGTATAAGAGAUAG UTGTUTUTTATAUAUATU TTGTTAATAAAGTGATTGTAGGTTGTA 3 *TCT****CCC TATAGT GAGTCGTATTA****TATG*

The other strand：

ATGTTGGATGTTAGTGAAATAATTGTTTUTAUAUATATTUTUTGTU GAUAGAGAATATGTGTAGAAAUAATTATT TUAUTAAUATUUAAUAT 5

5 CATA**TAATACGACTCACTATAGGG**AGA TACAACCTACAATCACT 。。。。。。。。。。

*GTAT****ATTATGCTGAGTGATATCCC*** *TCT* ATGTTGGATGTTAGTGAAATAATTGTTTUTAUAUATATTUTUTGTU GAUAGAGAATATGTGTAGAAAUAATTATT TUAUTAAUATUUAAUAT 5

5 CATA**TAATACGACTCACTATAGGG**AGATACAACCTACAATCACT *TTATTAACAAAAATATATATAAAAAACAA*………………………*CTATC TC TTATACA CATCT TTATTAATAA AATAATT AT AAATT ATA*

5. T7 transcription amplification:

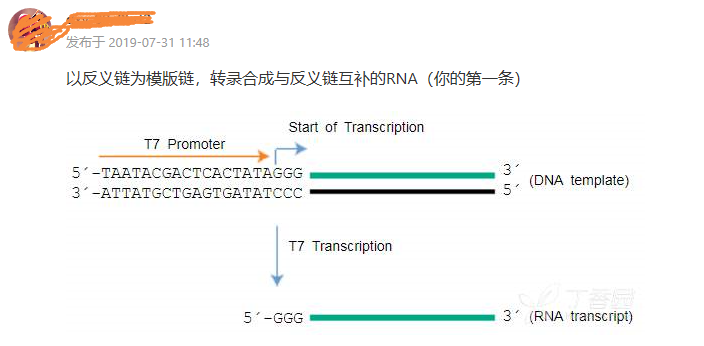

the RNA produced from T7 transcription:

5- GGGAGA UACAACC UACAAUCACU UUAUUAACAA AAAUAUAUAUAAAAAACAA………………………..CUAUC UC UUAUACA CAUCU *UUAUUAAUAA* AAUAAUU AU AAAUU AUA

6. Reverse transcription (Purple-reverse transcription primer):

5- GGGAGA UACAACC UACAAUCACU UUAUUAACAAAAAUAUAUAUAAAAAACAA………………………CUAUCUCUUAUACACAUCU*UUAUUAAUAA* AAUAAUU AU AAAUU AUA

CCC TC T ATGTTGG ATGTT A GTGA AAT AATT GTT TTT ATA TA TA TTT TTT GTT …………………… GAT AG AGAATATGTGTAGA

CATGAGACGCAACTAT GGTGACGAAUGATCA反转录引物

7. PCR amplification:

Template:

CCC TCTATGTTGG ATGTTAGTGA AATAATTGTT TTT ATATA TA TTT TTT GTT……………………….GATAGAGAATATGTGTAGA CATGAGACGCAACTAT GGTGACGAAUGATCA

First cycle:

AATGATACGGCGACCACCGAGATCTACACGGGAGATACAACCTA CAATCACT	09-P5 primer

CCCTCTATGTTGGATGTTAGTGAAATAATTGTTTTTATATATATTT TTTGTT ……….GATAGAGAATATGTGTAGACATGAGACGCAACTATGGTGACGAAUGATCA

5-AATGATACGGCGACCACCGAGATCTACACGGGAGATACAACCTACAATCACTTTATTAACAAAAATATATATAAAAAACAA------ -----CTATCTCTTATACACATCTGTACTCTGCGTTGATACCACTGCTTACTAGT-3

Second cycle:

AATGATACGGCGACCACCGAGATCTACACGGGAGATACAACCTACAATCACTTTATTAACAAAAATATATATAAAAAACAA----------CTATCTC TT ATACACATCTGTAC TCTGCGTTGATACCACTGCTTACTAGT-3

GATAGAGAATATGTGTAGACATGAG	09-P7 primer

TAGAGCATACGGCAGAAGACGAAC

Final PCR product：

AATGATACGGCGACCACCGAGATCTACACGGGAGATACAACCTACAATCACTTTATTAACAAAAATATATATAAAAAACAA----- CTATCTCTT ATACACAT CTGTAC TCATCT CGTATGCCG TCTT CT GCTTG

TTACTATGCCGCTGGTGGCTCTAGATGTGCCCTCTATGTTGGATGTTAGTGAAATAA TT GTT TTTATATATATT TT TTGTT -----GATAGAGAATATGTGTAGACATGAGTAGAGCATACGGCAGAAGACGAAC

### C. Protocol (TEAM-seq)

1. Adaptor anneal

5 CTGTCTCTTATACACATCT 3

5 TACAACCTACAATCACT NNNNNNNNNN AGATGTGTATAAGAGACAG 3

98℃ 1min； 30℃ 5min； 16℃	1h；

Agilent 2100 test the adaptor.

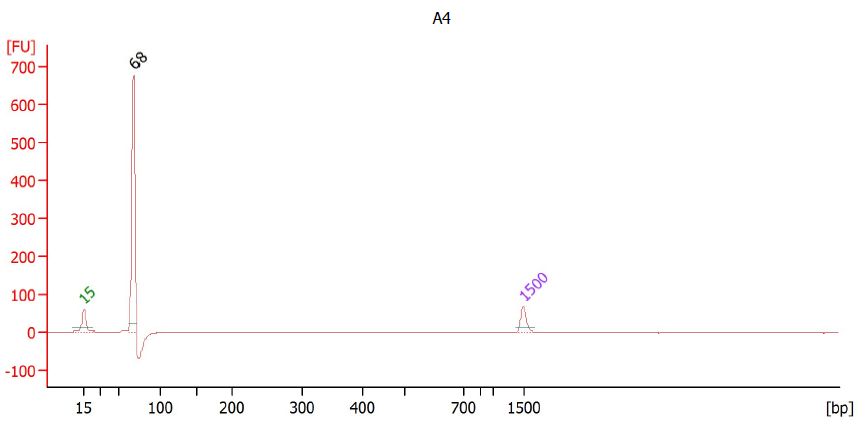

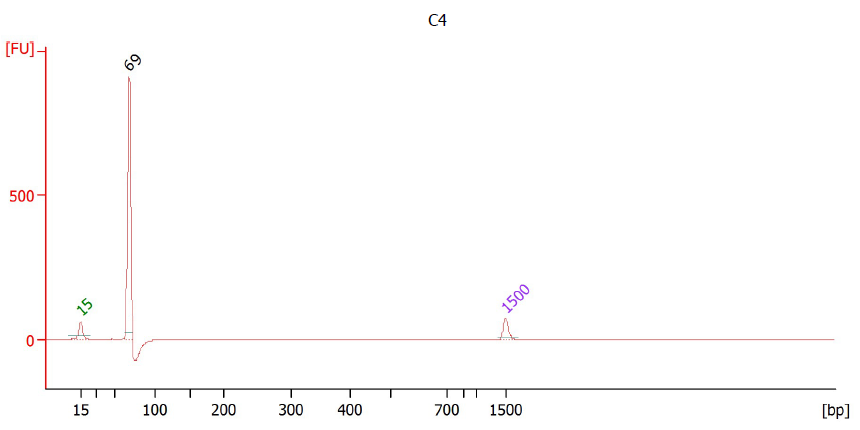

Transposon assembly

| 10uM Adapter | 2 |
| --- | --- |
| Tn5 （MGI） | 8 |
| Total | 10ul |
| 25℃ 1h（No heat cover） |  |

Tn5 tagmentation

| DNA+water | 7.5 | 37℃ 30min； |
| --- | --- | --- |
| 5Xtag H buffer（vazyme） | 2 |  |
| Tn5 | 0.5 |  |
|  | 10uL |  |

+1 uL 132mM EDTA (inactivated the Tn5)，65℃	20min.

+1 uL 216mM MgCl_2_ (neutralized the EDTA) ，RT 5min；

2. Gap filling

| DNA-biotin | 12 | 72℃ 10min； |
| --- | --- | --- |
| 5X Q5 buffer | 10 | 4℃ ∞ |
| 10mM each dNTP | 1 |  |
| Q5 DNA polymerase | 1 |  |
| NF water | 26 |  |
|  | 50 µL |  |

20ul M-280 Streptavidin beads（Invitrogen™ 11205D） binding for one hour (streptavidin protocol)；

3. EM-Seq Conversion (NEB E7125S)

| Reagent | Volume | |
| --- | --- | --- |
| DNA on beads | 28µl-bio | |
| 5X TET2 Buffer Supplement | 10 µl | |
| Oxidation Supplement | 1 µl | |
| DTT | 1 μl | |
| Oxidation Enhancer | 1 µl | |
| TET2 | | 4 µl |

| DNA (from step 2.2.) | 45 μl | 37℃ 1h ； |
| --- | --- | --- |
| Diluted Fe(II) Solution (adding 1 μl to 1249 μl of water.) | 5 µl |  |
| Total Volume | 50 µl |  |

Transfer the samples to ice and add 1 µl of Stop Reagent；Incubate at 37°C for 30 minutes

M280 magnetic stand pull down.

Denature:

| DNA on beads | 16 μl | 50℃ 10min；  Immediately place on ice |
| --- | --- | --- |
| 0.1 N NaOH | 4 µl |  |
| Total | 20µl |  |

Deamination

| The reaction from denatures | 20µl | 37℃ 3h； |
| --- | --- | --- |
| Nuclease-free water | 68µl |  |
| APOBEC Reaction Buffer | 10µl |  |
| BSA | 1 |  |
| APOBEC | 1 |  |
| Total volume | 100µl |  |

Place on a magnetic rack for 2minute.

—uses 1X AMPure XP beads to purify the supernatant ；add 10ul NF water to dissolve DNA;

—Wash the M280 beads 2 times….add 5ul NF water to resuspend magnetic beads；

—mix

4. Template extension

| T7 primer | 0.7 | 95℃ 1min |
| --- | --- | --- |
| DNA after EM-conversion | 16 | 65℃ 1min |
| 10X Phi 29 buffer | 2 | 40℃ 1min  4℃ ∞ |
| 2.5mM dNTP | 0.7 | 30℃ 20min |
| Phi29 | 0.6 | 65℃ 10min；  4℃ ∞；  转移冰上至少5min； |
|  | 20uL |  |

5. T7 transcription (NEB E2040S)

| The reaction from last step | 20 | 37℃ 18h； |
| --- | --- | --- |
| 10X T7 buffer | 6 | 4℃ ∞ |
| NTP | 10 |  |
| T7 RNA polymerase | 2.5（3.5） | 中途补加1ul T7酶； |
| RNase inhibitor | 1 |  |
| NF water | 20.5 |  |
|  | 60uL |  |

1.8X RNA Clean XP purification，12ul NF water elution；

Agilent 2100 RNA pico

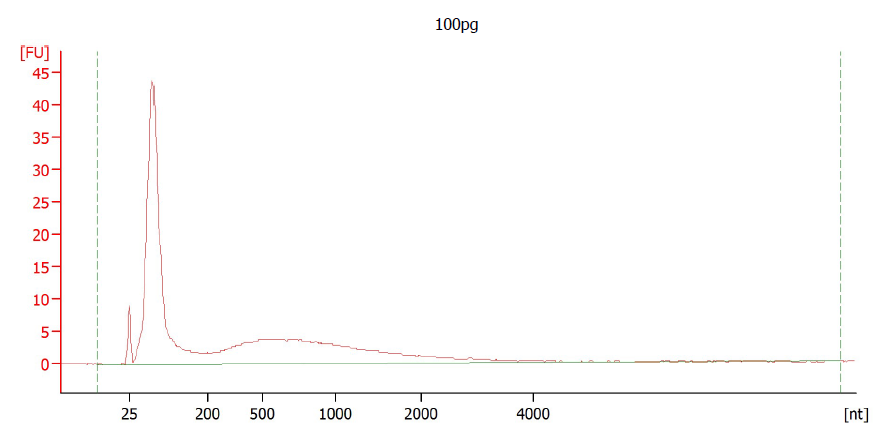

6. Reverse transcription (Invitrogen 18090050)

| 10uM RT primer | 1 | 65℃ 5min，then on ice； |
| --- | --- | --- |
| RNA template from last step | 10 |  |
| 5X SSIV buffer | 2.5 |  |
|  | 13.5 |  |

| 5X SSIV buffer | 1.5 | 55℃ 15min |
| --- | --- | --- |
| 0.1 M DTT | 1 | 60℃ 10min |
| 10mM dNTP | 1.5 | 65℃ 12min |
| RNase inhibitor | 1 | 70℃ 8min |
| SSIV | 1 | 75℃ 5min |
|  | 6.5 | 80℃ 10min；  4℃ ∞ |

7. PCR amplification（Roche KK2801）

| 2X KAPA Uracil mix | 25 | 95℃ 3min |
| --- | --- | --- |
| RT primer/PCR primer | 1.2（0.6+0.6） | 98℃ 20s |
| cDNA | 7 | 60℃ 20s 11cycle |
| NF water | 16.8 | 72℃ 4min |
|  | 50uL | 72℃ 10min；  4℃ ∞ |

0.6X Ampure XP purification.

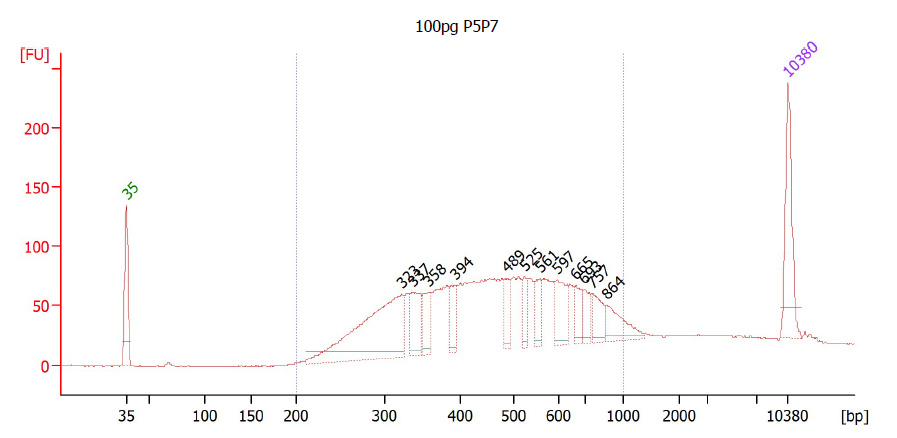

Single cell indexed protocol

1. **Nuclei isolation**

Refer to 10Xgenomics protocol “nuclei isolation for single cell ATAC sequencing” or other suitable nuclei isolation protocol for your sample; resuspended in PBS and cell sorting (BD cell sorting protocol)

2. **Cell lysis &Tn5 tagmentation**

| Cell（nuclei）+water | 2.5 | 55℃ 1h； |
| --- | --- | --- |
| Tris-Cl（pH8.0）120mM | 1 （30mM） | 70℃ 20min |
| Qiagen Protease (4mg/ml) | 0.5 |  |
|  | 4 uL |  |
| Cell lysis mix | 4 | 37℃ 30min |
| 5Xtag H buffer（vazyme） | 1.4 |  |
| Tn5（30Xdilution） | 0.6 |  |
|  | 6uL |  |

+0.5 uL 156mM EDTA (final concentration 12mM)，65℃	20min.

+0.5 uL 252mM MgCl_2_ (final concentration 18mM) ，RT 5min；

After the tagmentation, the cells could be pooled and processed together.

3. **Gap filling**

| DNA-bio | 7 | 72℃ 15min； |
| --- | --- | --- |
| 5X Q5 buffer | 2.4 | 4℃ ∞ |
| 10mM each dNTP | 0.7 |  |
| Q5 DNA polymerase | 0.7 |  |
| NF water | 1.2 |  |
|  | 12 µL |  |

Dynabeads™ M-280 Streptavidin to bind the DNA fragments (M280 protocol).

### D. Other methods (MDA, Trueprime, PEG+TMAC, BST)

Other methods showed in Figure2, included the EM-MDA-seq, EM-MDA-seq(Trueprime), EM-MDA-seq with addictives, EM-MDA-seq with BST, EM-MDA-eq with GpC pretreated. The genomic DNAs were directly treated with EM-seq kit (NEB E7120S) as EM-seq protocol. The converted single stranded DNAs (~10kb) were purified by 1X Ampure xp beads (A63881). Then the DNAs underwent the MDA amplification by REPLI-g Single Cell Kit (Qiagen 150343). For Trueprime MDA-EM-seq, the genomic DNAs underwent the protocol SYGNIS TruePrimeTM WGA & Single Cell WGA Kits (Lucigen), instead of the REPLI-g single cell kit. In the EM-MDA-seq with additives, all the addictive(3% PEG,100mM TMAC, 3%PEG+100mM TMAC) were added to the REPLI-g Single Cell reaction. In the EM-MDA-seq with BST amplification, we used the BST2.0 (NEB M0537S) to substitute the Phi29 in REPLI-g single cell kit.

[1] Zerbino DR, Wilder SP, Johnson N, Juettemann T, Flicek PR. The ensembl regulatory build. Genome Biology. 2015 Mar;16:56. DOI: 10.1186/s13059-015-0621-5. PMID: 25887522; PMCID: PMC4407537.

[2] Smallwood, S., Lee, H., Angermueller, C. *et al.* Single-cell genome-wide bisulfite sequencing for assessing epigenetic heterogeneity. *Nat Methods* 11, 817–820 (2014). https://doi.org/10.1038/nmeth.3035

[3] ENCODE Project Consortium. An integrated encyclopedia of DNA elements in the human genome. Nature. 2012 Sep 6;489(7414):57-74. doi: 10.1038/nature11247. PMID: 22955616; PMCID: PMC3439153.
